# Modelling cell adaptation using internal variables: accounting for cell plasticity in continuum mathematical biology

**DOI:** 10.1101/2023.05.08.539423

**Authors:** Marina Pérez-Aliacar, Jacobo Ayensa-Jiménez, Manuel Doblaré

## Abstract

Cellular adaptation is the ability of cells to change in response to different stimuli and environmental conditions. It occurs via phenotypic plasticity, that is, changes in gene expression derived from changes in the physiological environment. This phenomenon is important in many biological processes, in particular in cancer evolution and its treatment. Therefore, it is crucial to understand the mechanisms behind it. Specifically, the emergence of the cancer stem cell phenotype, showing enhanced proliferation and invasion rates, is an essential process in tumour progression.

We present a mathematical framework to simulate phenotypic heterogeneity in different cell populations as a result of their interaction with chemical species in their microenvironment, through a continuum model using the well-known concept of internal variables to model cell phenotype. The resulting model, derived from conservation laws, incorporates the relationship between the phenotype and the history of the stimuli to which cells have been subjected, together with the inheritance of that phenotype. To illustrate the model capabilities, it is particularised for glioblastoma adaptation to hypoxia. A parametric analysis is carried out to investigate the impact of each model parameter regulating cellular adaptation, showing that it permits reproducing different trends reported in the scientific literature. The framework can be easily adapted to any particular problem of cell plasticity, with the main limitation of having enough cells to allow working with continuum variables. With appropriate calibration and validation, it could be useful for exploring the underlying processes of cellular adaptation, as well as for proposing favorable/unfavourable conditions or treatments.

## 1 Introduction

Cellular adaptation comprises the changes induced in cell behaviour in response to changes in their environment. It can occur via natural selection (long term response) or via phenotypic plasticity, which is the ability of a cell with a given genotype to produce different phenotypes in reaction to environmental changes [1]. There are many examples of phenotypic plasticity in nature. For example, some butterflies develop different wing colours depending on factors such as the amount of food available [2]. Another strategy is bet-hedging, consisting in randomly diversifying into different populations so that, should the environment change, some of them would have a higher probability of surviving [3]. Cells can also be reprogrammed towards stem-like phenotypes and then differentiated to other cell types [4, 5].

Epigenetics is the study of heritable changes in gene expression (phenotype) which are not caused by alterations in the DNA sequence or genotype (detailed information about epigenetic mechanisms can be found in [6, 7]). Epigenetic changes can be triggered by the environment and hence, they are key in cell plasticity [7]. In a sense, epigenetics is the link between the environment and phenotypic plasticity leading to cellular adaptation [8]. Epigenetic mechanisms are responsible for cell differentiation, since all cells in our organism have the same genotype, and it is their different gene expression what characterises them [9]. Throughout our lifespan, we are subject to epigenetic changes depending on environmental factors, such as diet, habits (smoking) and social interactions [10]. Besides, epigenetics is involved in the occurrence and progression of diverse human diseases [11, 12] and, in particular, in cancer. While cancer was initially thought to be caused by the accumulation of genetic mutations, in the past decades, epigenetics have been shown to play a central role in cancer development and progression [13, 14].

Tumours are now recognized as heterogeneous cell populations, presenting both genetic and non-genetic (epigenetic) differences among them [15]. Indeed, in 2022, phenotypic plasticity has been included among cancer hallmarks, and epigenetic reprogramming as an enabling characteristic facilitating the acquisition of this hallmark capability [16]. Phenotypic plasticity explains some of the most characteristic features of cancer, such as metastasis and drug resistance, probably the main challenges for improving cancer prognosis [17]. Cells, in order to become metastatic, must acquire some plasticity following the Epithelial to Mesenchymal Transition (EMT), by which cells cells increase their motility [18]. This transition is triggered by different environmental factors, such as epigenetic reprogramming or external factors like hypoxia, which are now object of important research [19]. Among the different populations present in tumours, Cancer Stem Cells (CSCs) have gained much attention in recent years [20]. These cells are believed to drive tumour initiation and growth, as well as to be related with aggressiveness and therapy resistance [21]. In opposition to differentiated tumour cells, CSCs share capacities with normal stem cells, having a higher capacity of self-renewal. The tumour population is in permanent evolution, with cells moving from differentiated phenotypes to stem-like phenotypes and vice versa depending on the Tumour MicroEnvironment (TME) [22]. Also, the EMT has been closely related to the acquisition of stem phenotypes [23].

Hypoxia is one of the main features of solid tumours, due to their high oxygen demand and their aberrant vasculature [24]. Hypoxic tumours usually correlate with increased aggressiveness and drug resistance. Current evidence supports that hypoxia drives cells towards CSC phenotypes, whereas high oxygen levels promote differentiation [25, 26]. Thus, the effects that hypoxia causes in tumours and the mechanisms behind their enhanced aggressiveness are currently a hot topic in cancer research [27]. In this work, we take the example of glioblastoma (GBM) evolution under hypoxic conditions. GBM is the most common and lethal primary brain cancer, with a 5-year survival rate of only 6.8% [28], rendering it one of the cancers with worst prognosis. Hypoxia is a defining feature of GBM, and a major concern for its prognosis, since it increases cell aggressiveness, increasing the cells capacity to proliferate and invade the surrounding tissue [29]. Hence, it is important to better understand how hypoxia triggers cellular adaptation in GBM, and in tumours in general, since it can help improving cancer prognosis and treatment response.

Mathematical models are valuable tools to understand complex phenomena relating tumours and their TME and test hypotheses regarding the effects of different environmental conditions on the adaptive response of tumours. In the last years, some mathematical models have been developed to deal with cellular phenotypic plasticity and adaptation from different perspectives, ranging from agent-based models [30, 31, 32] to continuum ones [33, 34, 35, 36]. Here we focus on the latter type. This family constitutes the majority of adaptation models developed to date and they use well known formalisms as partial integro-differential equations which are easily implemented and allow modelling the most relevant phenomena in cancer evolution together with the interactions with the TME. Several of these models define a finite number of phenotypes that behave differently, with the transitions between them mediated by environmental conditions. This approach has been widely applied to model drug resistance [37, 38, 39], but also to other adaptation processes such as hypoxia-driven adaptation [35]. However, this discrete approach to the phenotypic state does not correspond with biological observations, in which cells go through a wide spectrum of different phenotypes caused by epigenetic changes [34]. Hence, it makes more sense to consider gene expression as a continuum variable. Very recently, some authors have followed this standpoint, defining a new *artificial* dimension corresponding to the cell’s phenotypic state. Some of these models consider adaptation as driven by random effects (genetic mutations) [33], while others incorporate the effect of the environment as a key agent driving phenotypic evolution [34, 36].

In this work, we propose a new mathematical framework to model cell adaptation processes and phenotypic plasticity driven by the environment. The model is focused on tumour evolution, but its formulation is general and could be particularised to any cell population and/or biological process. It considers cell phenotype as a continuum variable, but instead of adding an extra dimension to the problem [34, 36], the phenotype is modelled as a set of internal variables that define the cell state, and consequently the cells response. This allows an easy physical interpretation of the internal variables and their relationship with the environment, while keeping the idea of continuum phenotype. The proposed approach closely follows the concept of *state* from control systems or non-linear mechanics. Cell phenotype (or state) represents at the cell level the changes at the molecular level (e.g. epigenetic changes) that lead to alterations in the cell’s gene expression and therefore in its expression. A differential equation, analogous to the ones defined for cells and chemical species, is derived to describe the evolution of each internal variable, relating their change with different signal levels coming from the TME. The activation functions regulating cell behaviour are now also dependent on those internal variables. Thus, the proposed framework allows to incorporate cell response to environmental changes as well as the reversibility and inheritance typical of phenotypic changes.

The framework is first formulated in Section 2, for an arbitrary number of cell populations, chemical species and internal variables. Then, we particularise it to the case of GBM evolution under hypoxic conditions, the example chosen to illustrate the model capabilities. In Section 3, an extensive parametric analysis is performed to analyse the effect of the different parameters regulating cell state evolution and to show the potential of the model for capturing different trends. Next, in Section 4, we study GBM evolution under different environmental conditions. In Section 5, we discuss the capabilities and the results of our approach to situate it within the existing approaches to model cell adaptation in literature. Finally, Section 6 presents the main conclusions of the work.

## 2 Mathematical framework

In this section, we present the general mathematical framework developed to describe cell adaptation processes and phenotypic heterogeneity in tumour evolution. This framework can then be particularised to simulate a wide range of different problems, depending on the precise conditions and parameters, although in this paper, we focus on glioblastoma evolution under hypoxic conditions.

### 2.1 General framework

The starting point is a previously published continuum model [40, 41] describing the spatio-temporal evolution of different cell populations and chemical species constituting the TME. In what follows, *T* and *X* are used to represent the temporal and spatial coordinates respectively. All variables are considered at the population level, within a continuum framework. Considering the concentration of *m* cell populations *C*_*i*_ = *C*_*i*_(*X, T*) (*i* = 1, ⋯, *m*) and *n* chemical species *S*_*i*_ = *S*_*i*_(*X, T*) (*i* = 1, ⋯, *n*), we can write a vector of field solutions ***U*** with *m* + *n* components, formed as:

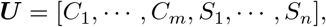

The transport equation for each component *U*_*i*_ is then written as:

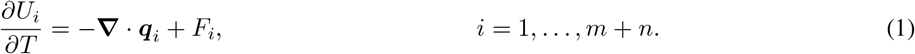

In Eq. (1), ***q***_*i*_ is the flux term, which in general can include both diffusive and convective terms. Analogously, *F*_*i*_ is the source term. In the case of cell populations, the source term *F*_*i*_ comprises proliferation and death:

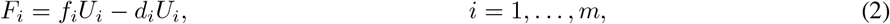

with *f*_*i*_ the growth rate and *d*_*i*_ the death rate.

For the chemical species, the source term includes the phenomena of decay, as well as production and/or consumption by cells. We can therefore write these terms as:

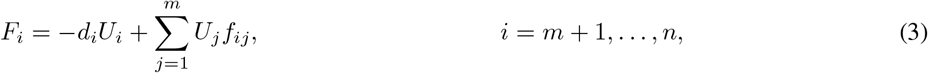

with *d*_*i*_ the decay rate, and *f*_*ij*_ the function modelling the consumption or production rate of the *i*^th^ species by the *j*^th^cell population. In general, all the aforementioned terms and functions (***q***_*i*_, *f*_*i*_, *d*_*i*_, *f*_*ij*_) may depend on any solution field and their derivatives. That is, every phenomena involved in cell evolution can depend on the current cell and species concentrations and on the changes in these quantities. Their particular functional form will depend on the problem in hands, and will be defined later for our reference problem, GBM evolution.

Once this primary model has been defined, we extend it to incorporate the phenomenon of cell adaptation, taking into account the cell history and how they keep “in memory” their past states. To do that, we introduce the concept of cell state (or phenotypic state) [42], which is described by one or more internal variables, that jointly represent the current phenotypic state of the cell. This is based on the state theory, widely used in other disciplines such as control theory [43] or non-linear mechanics [44] as a way to model phenomena that are dependent on past states. These variables constitute a macroscopic representation of the microscopic changes inside a cell that lead to phenotypic heterogeneity, for instance, epigenetic changes [7, 45].

In general, we consider that the state of each cell population *i* can be fully described by *r*_*i*_ internal variables, accounting for the cell phenotypic changes in response to different intrinsic or extrinsic stimuli. Each variable must lie within its corresponding state space, which comprises the set of all possible configurations. Let *V*_*ik*_ denote the *k*^th^ internal variable affecting the cell population *U*_*i*_, for *i* = 1, ⋯, *m*, with 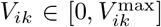. We can write the corresponding evolution equation as:

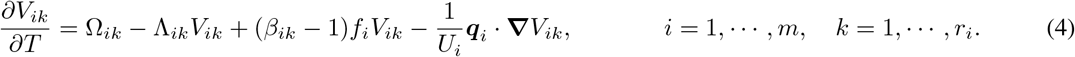

Since this is a population model, internal variables are measured in the corresponding units per unit cell, expressing the average epigenetic changes per cell in the population of a Representative Volume Element (RVE) that lead to the current phenotypic state, as usual in continuum physics. The details of the mathematical derivation of Eq. (4), from the perspective of conservation laws, can be found in Appendix **??**. The meaning of the different terms in Eq. (4) is detailed next:

- In the first term, Ω_*ik*_ represents the function of epigenetic changes acquisition and may depend on a combination of external stimuli, their derivatives, and the current level of the internal variables themselves. This allows to model cell adaptation as a response to environmental changes.
- The second term in Eq. (4) is the decay term, with Λ_*ik*_ the decay coefficient. This term accounts for the natural reparation paths that cells follow to overcome genetic and epigenetic mutations, which are, in general, reversible [46, 47, 48].
- The third term takes into account the fact that epigenetic changes may be inherited through cell division [49, 50, 51]. In this regard, the term is proportional to the growth rate in cells (*f*_*i*_) through the coefficient *β*_*ik*_. If *β*_*ik*_ = 1, daughter cells inherit the same phenotypic state as their progenitor, whereas if *β*_*ik*_ < 1, only a percentage of daughter cells inherits the state of the parents. Otherwise, this could be interpreted as a partial repair of the phenotypic changes present in those parent cells.
- Finally, the last term is a convection term, necessary to adapt the state equation to our Eulerian framework (see Appendix **??** for the details). Each internal variable *V*_*ik*_ is convected with the flux term ***q***_*i*_ of its related cell population *U*_*i*_.

Lastly, we must define the effect that internal variables may have on cell behaviour. In the most general case, the set of internal variables ***V*** _*i*_ = (*V*_*ik*_), *k* = 1, …, *r*_*i*_, associated with the population *U*_*i*_, (for *i* = 1, ⋯, *m*, since we assume that internal variables affect cell behaviour, but not chemical species) could affect every mechanism involved in the evolution of this population, so we would have:

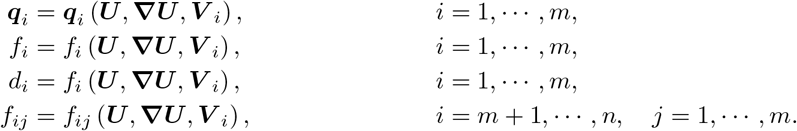

### 2.2 One-dimensional model of GBM evolution under hypoxic conditions

Next, we particularise the proposed general model to a specific problem, namely the evolution of GBM subjected to oxygen variations. For the sake of simplicity in the implementation and computations, we shall consider a one-dimensional model with two cell populations, corresponding to alive (*C*_a_) and dead (*C*_d_) cells respectively, a chemical species which is oxygen (*S*) and an internal variable (*V*), accounting for the effects of hypoxia on the phenotypic state of GBM cells.

This model is based on a previous work [52], which is extended by incorporating this internal variable, thus allowing for the simulation of more complex processes such as those of cell memory and adaptation. The reader is therefore referred to [52] for further details about the model and the physical meaning of its parameters and functions.

The equations governing the evolution of the concentration of alive and dead cells are:

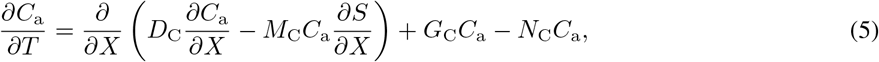

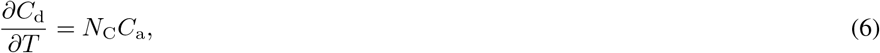

where *D*_C_ is the diffusion term and *M*_C_ is the chemotaxis term. They jointly model cell movement, both in a random way (diffusion or pedesis) and towards oxygen gradients (chemotaxis). With regards to the source term, it comprises cell proliferation and death through the terms *G*_C_ and *N*_C_ respectively.

The corresponding coefficients may be written as:

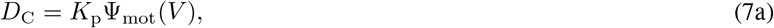

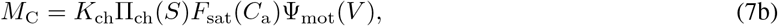

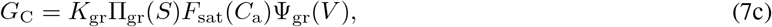

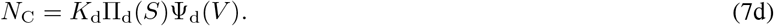

As can be seen in the above equations, all terms may be affected by the phenotypic state through the Ψ functions, which will be detailed later, as we define more precisely the internal variable *V*, its meaning and the hypotheses related to its effects on cell behaviour.

The diffusion or pedesis term (Eq. 7a) is considered constant (except for the influence of phenotypic changes), with *K*_p_ the pedesis coefficient.

Both chemotaxis (Eq. 7b) and proliferation (Eq. 7c) are regulated by the *go or grow* hypothesis [53] in GBM evolution, which states that cells spend their resources either in proliferating or migrating, depending on the oxygen level and, in particular, on the hypoxia threshold *S*^*H*^. To model this dependence with oxygen, we use for both phenomena the ReLU-like activation functions Π_ch_, Π_gr_ [52]:

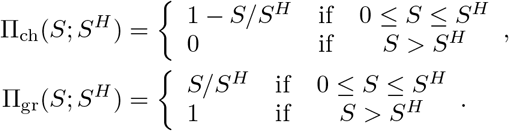

Also, both phenomena are restricted by space considerations assuming that cells cannot proliferate nor migrate in saturated areas [54, 52] by means of a logistic growth model. Hence, we define *F*_sat_ to take into account this effect:

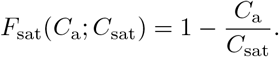

As for the coefficients regulating these phenomena, *K*_ch_ is the chemotaxis coefficient, while *K*_gr_ is the characteristic growth rate.

Cell death (Eq. 7d) is regulated by *K*_d_, the characteristic death rate, also depending on the oxygen concentration, with cells mainly dying below an anoxia threshold. This is modelled with an hyperbolic tangent activation function, depending both on a location parameter *S*^*A*^ and a spread parameter Δ*S*^*A*^, to allow stochastic death (apoptosis) as well [52]:

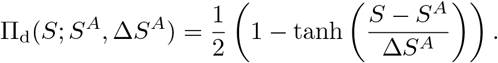

Moving on to chemical species, the governing equation for the oxygen concentration is:

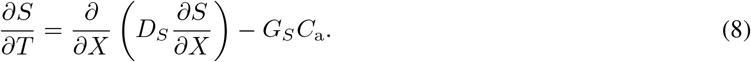

As can be seen in Eq. (8), the evolution of oxygen is driven by diffusion through the diffusion term *D*_*S*_ and by a source term accounting for cell oxygen uptake *G*_*S*_. These terms are written as:

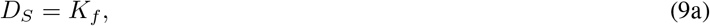

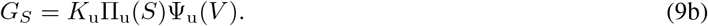

As for the cells, the oxygen diffusion term (Eq. 9a) is constant, with *K*_*f*_ the diffusion coefficient. Oxygen uptake by alive cells (Eq. 9b) is regulated by the uptake rate *K*_u_ and Π_u_, a nonlinear correction function accounting for the dependence between cell uptake and the oxygen level, following oxygen consumption kinetics [55]:

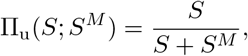

with *S*^*M*^ the Michaelis-Menten constant. The value of the parameters regarding GBM evolution have been obtained from [52], where they were fitted to different *in vitro* experiments in microfluidic devices.

Finally, we define an internal variable *V* which models the effects of hypoxia on cell state. GBM cells are known to undergo phenotypic changes under hypoxia, which promote their dedifferentiation towards a cancer stem cell like phenotype [56]. The variable *V* represents the state of the cell (comprising factors such as gene expression levels or the number and position of DNA methylation) in an averaged macroscopic sense in the RVE population. It is bounded, so *V* ∈ [0, *V*_max_]. When *V* = 0, the state of the cell corresponds to a totally differentiated GBM cell, while for *V* = *V*_max_, the cell behaves like a cancer stem cell. The governing equation for the evolution of the internal variable is phenomenologically expressed as:

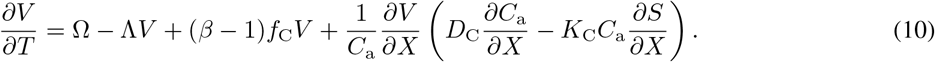

In absence of suitable experimental data, it is difficult to accurately define the functional forms as well as the value of the parameters involved in Eq. (10). Hence, in what follows we propose reasonable functional forms and parameters which may help us to show the potential of the model for qualitatively reproducing biologically-consistent behaviours, in compliance with what has been reported in literature.

We consider that the phenotypic changes are driven by external factors, in particular by the oxygen level, such that cells go towards a more stem like phenotype when the oxygen level is below the hypoxic threshold *S*^*H*^ while cells may differentiate again when the oxygen level is above *S*^*H*^. To model this, and taking into account that *V* is bounded, we define Ω as:

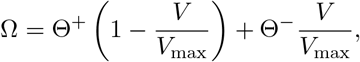

with Θ^+^ = max(0, Θ), Θ^−^ = min(−Θ, 0) and the function Θ defined as:

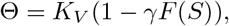

where *K*_*V*_ is the rate at which epigenetic changes are acquired (i.e., the rate at which the phenotypic state is modified) and *γ* ∈ [1, 2] is the parameter regulating whether cells move towards a differentiated phenotype when *S > S*^*H*^ (*γ* = 2) or not (*γ* = 1), that is, whether the epigenetic change is elastic/inelastic. *F* is an asymmetric generalised normal distribution function, *F* (*S*) = Φ(*y*), with Φ the normal cumulative distribution function and *y* is defined as:

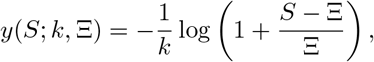

with *k* < 0 the shape parameter and Ξ > 0 the location parameter. This function is convenient since it has positive support (the oxygen level cannot be negative) and allows enough flexibility to consider different dependencies and the possibility (or not) of phenotype changes above *S*^*H*^ in a non-symmetric way.

The second, third and fourth terms in Eq. (10) correspond to the decay, heritage and convection of the internal variable, respectively, as explained in Section 2.1, with Λ the decay rate and *β* the degree of phenotypic heritage.

Regarding the effects of phenotypic changes in cell behaviour, we define the Ψ functions as beta distribution functions, which are bounded and therefore coherent with our definition of *V* as a bounded function. They also allow great flexibility for modelling different behaviours. In general, we write:

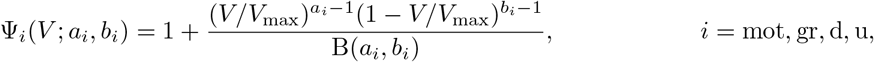

with:

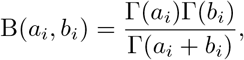

and Γ the Gamma function.

#### Model non-dimensionalisation

To simplify the interpretation of the *in silico* results, the model is reformulated in a dimensionless form, using the following dimensionless variables:

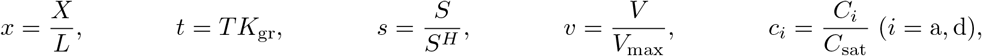

where *L* is the length of the computational domain, *L* = 0.2 cm. The value of the rest of the parameters used to define the dimensionless variables are *K*_gr_ = 200 h, *S*^*H*^ = 7 mmHg, *C*_sat_ = 5 10^7^ cell*/*mL (obtained from [52]) and *V*_max_ = 1 cell^*−*1^ (arbitrarily set to define the *V* value corresponding to the CSC phenotype).

With such definitions, Eqs. (5,6,8,10) can then be expressed as:

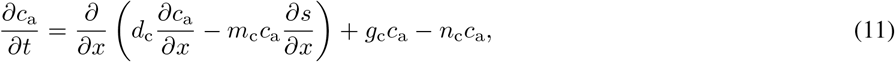

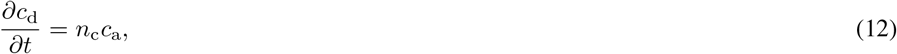

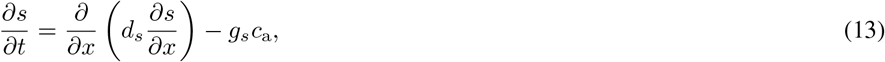

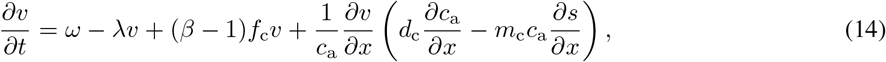

with

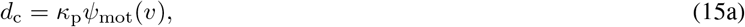

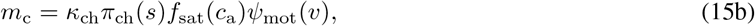

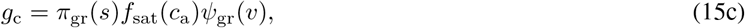

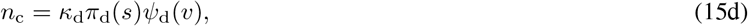

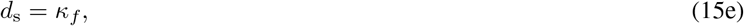

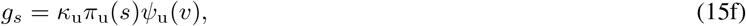

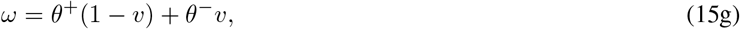

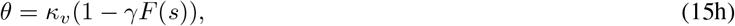

and the following dimensionless parameters:

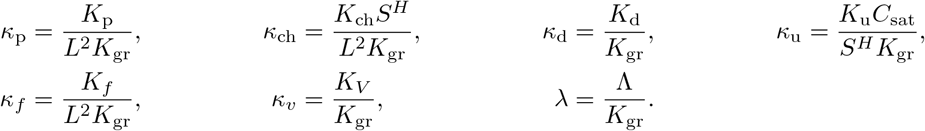

The non-dimensional activation functions are expressed as:

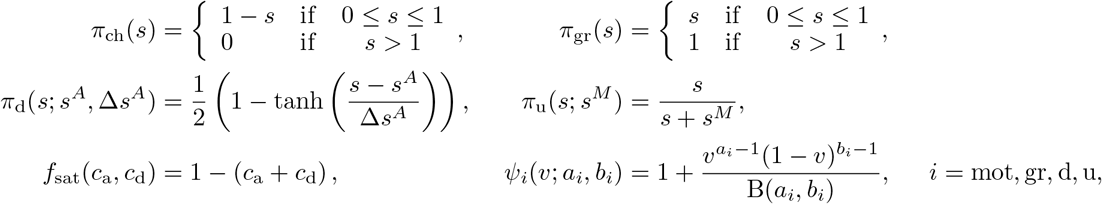

with the parameters:

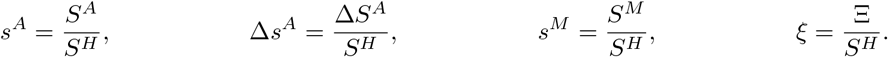

The presented model has to be completed with suitable initial and boundary conditions.

#### Model parameters

As stated before, the parameters related to cell evolution are taken from a previous model [52]. The corresponding non-dimensional values are included in Table 1.

**Table 1:** Dimensionless parameters related with cell evolution (adapted from [52]).

| Parameter | Value |
| --- | --- |
| $\kappa_p$ | $9.0 \cdot 10^{-3}$ |
| $\kappa_{ch}$ | 0.95 |
| $\kappa_d$ | 0.24 |
| $\kappa_u$ | $5.1 \cdot 10^3$ |
| $\kappa_f$ | $1.6 \cdot 10^3$ |
| $s^A$ | 0.23 |
| $\Delta s^A$ | $1.4 \cdot 10^{-2}$ |
| $s^M$ | 0.36 |

The value for the rest of the parameters is estimated within reasonable biological ranges and for illustrative purposes. In particular, the value of *κ*_*v*_ is set to 0.72 and the decay rate is assumed to be zero (*λ* = 0). The value of the parameters defining the function *θ* will be further explored in the next section. Regarding the parameters related to effect of the phenotype on cell behaviour, i.e., the parameters in the *ψ* functions, we define them to achieve functions consistent with biological evidence:

- Cells with a stem-phenotype have increased proliferation [57, 58, 59, 60].
- Cells with a stem-like phenotype undergo the EMT and hence, increase their migratory activity [57, 61].
- Stem cells are more resistant to apoptosis than differentiated cells and therefore, they present an overall lower death rate [62, 63].
- Due to their increase in metabolic activity, cells with a stem-like phenotype consume more resources, and thus have a higher uptake rate [64].

The precise values of the parameters as well as the shape of the functions are presented in Figure 1.

**Figure 1:**
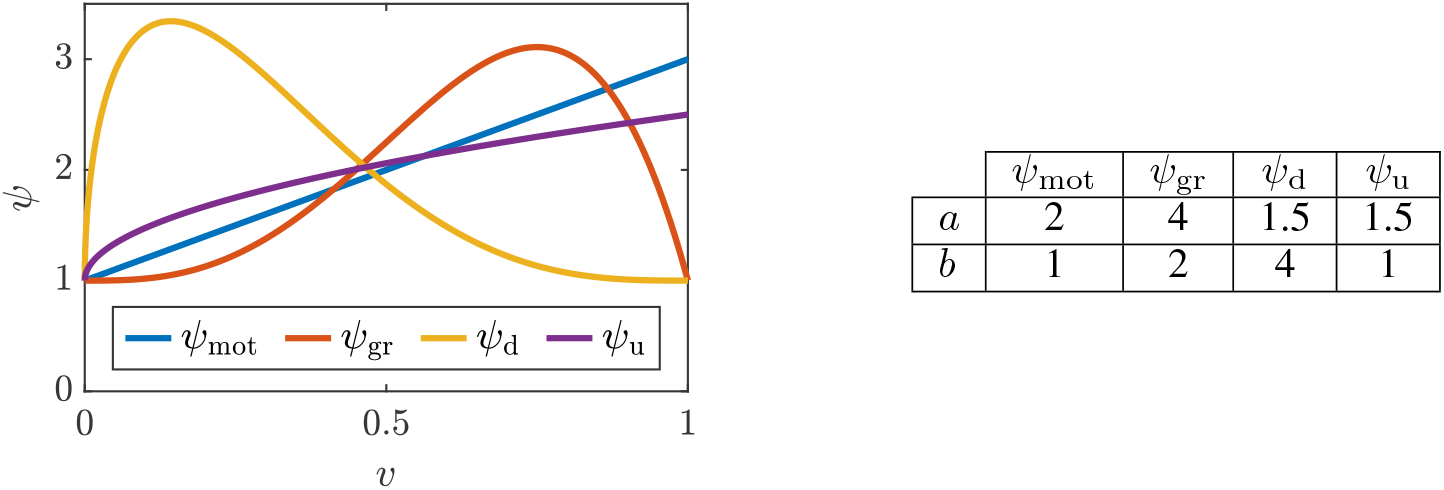
Shape of the *ψ* functions for the different phenomena, together with the value of the corresponding parameters for each function. *ψ*_mot_ and *ψ*_u_ are monotonically increasing functions; while *ψ*_gr_ and *ψ*_d_ are both non-monotonic, they increase, reach a maximum, and then decrease, reflecting that cells are most active in the phenotype corresponding to the maximum in the *ψ* function.

## 3 Model Inspection

### 3.1 Model set-up

In this section we investigate the effect of the different parameters involved in the evolution model of the internal variable (Eq. 14) in a benchmark experiment of GBM evolution under cyclic hypoxia in microfluidic devices. Microfluidic techniques allow nowadays to reproduce, in a controlled microenvironment, the three dimensional estructure of tumours. In a previous work [52], a model of GBM evolution under hypoxia based on experiments on microfluidic devices was developed. In this work we continue to embrace this framework, since the recent development and advances in microfluidics enable the generation of large amounts of data which may be used for validation purposes.

The experimental configuration is one-dimensional, and recreates cells within the central chamber of a microfluidic device, of width *L*, with two lateral channels through which oxygen-rich medium can be perfused. This configuration simulates cells between two blood vessels in the brain, and allows controlling oxygenation conditions and creating oxygen gradients within the chamber. To subject cells to cyclic hypoxia, we simulate alternative oxygen perfusion through the channels, in cycles of *T* = 7 days (*t* = 0.84). A scheme of the device configuration and our cyclic hypoxia *in silico* experiment are shown in Figure 2.

**Figure 2:**
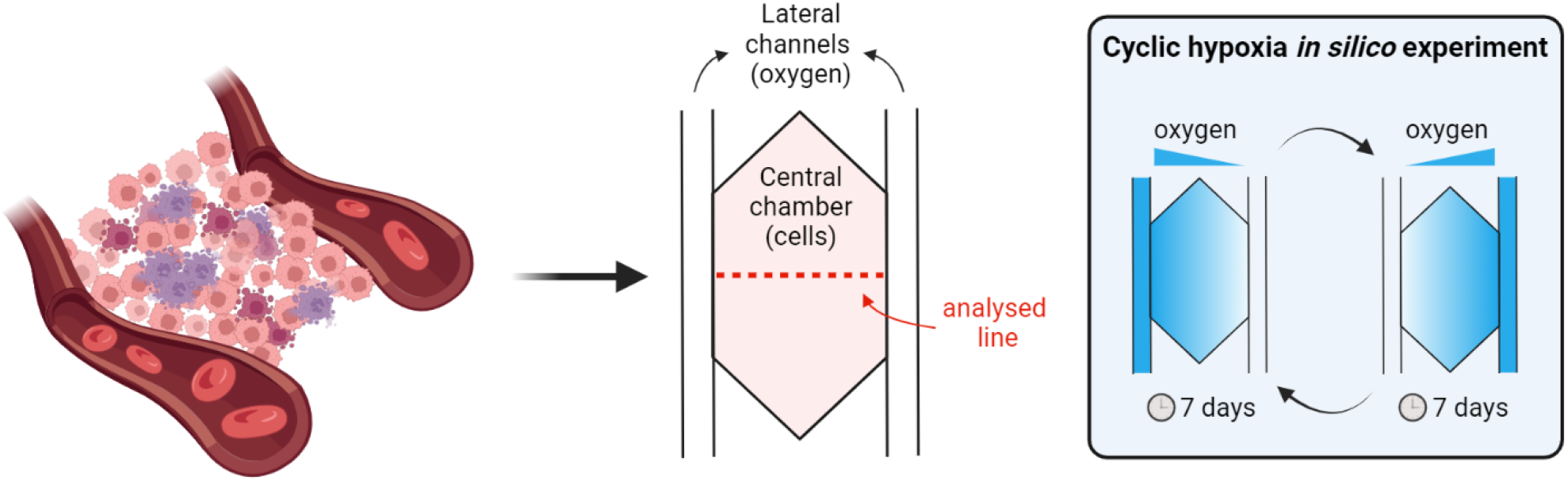
Schematic representation of the microfluidic device and the experiment modelled. The microfluidic device reproduces tumoural cells evolution between two blood vessels, which supply oxygen and nutrients. The experiment of cyclic hypoxia consists on perfusing oxygenated medium alternately through each channel while keeping the other one sealed, thus creating gradients. Created with Biorender.com

The level of oxygen is above the hypoxic threshold in the oxygenated channel, *s* = 9*/*7, and below it in the non-oxygenated channel, *s* = 1*/*7. Consequently, we impose time-dependent Dirichlet boundary conditions for oxygen in both channels, with the corresponding value at each time point. The initial condition is set to be a straight line joining both channels, with constant gradient between the two boundary conditions. This is due to the quick oxygen diffusion, which causes the stationary profile to be reached in a short period of time, compared with the characteristic time of the cell processes considered.

Regarding alive cells, we assume that they are initially uniformly distributed within the chamber, at a low concentration *c*_a_(*x, t* = 0) = 0.08 (far from the saturation limit). We assume that, initially, there are no dead cells (*c*_d_(*x, t* = 0) = 0). Neither alive nor dead cells can go away from the chamber, so homogeneous Neumann boundary conditions are imposed for cells. The initial phenotypic state is uniform *v*(*x, t* = 0) = *v*_0_ and, in order to explore its impact, it will be varied in each simulation.

We simulate 4 cycles of hypoxia, with a time step of Δ*t* = 0.0014 (equivalent to 1000 s) and a spatial discretisation with an element size of Δ*x* = 0.005 (equivalent to 10 *μ*m. The system of Partial Differential Equations (PDEs) is implemented in MATLAB and solved with the library pdepe [65].

To analyse the results, we introduce some metrics to represent the global macroscopic state of the tumour, aggregating spatial effects. First, we define the Tumour Burden (TB) as the total number of alive cells, as a measure of the size of the active tumour at each time:

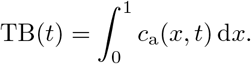

Analogously, we define the Mean phenotypic State (MS) as the mean phenotype in the population at each point and time:

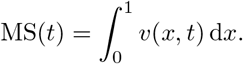

### 3.2 Parametric analysis

First, in Figure 3, we present a summary of simulations varying the parameters related with the evolution of the phenotypic state. Simulations have been performed and analysed for different values of the parameters that determine the function *F*, which establishes how the phenotypic state changes due to specific environmental conditions (in this case, due to the oxygen concentration). Also, we perform simulations for different values of the parameter *β*, which determines how cells inherit the phenotypic state of their progenitors. Finally, we have studied three different scenarios regarding the initial cell preconditioning, ranging from a tumour mainly composed of differentiated cells (*v*_0_ = 0.1) to a tumour predominantly formed of CSCs (*v*_0_ = 0.9). Details of these and other simulations, showing the whole temporal evolution of the Tumour Burden and Mean phenotypic State for the different parameters involved in the evolution of the phenotypic state, can be found in Appendix **??**.

**Figure 3:**
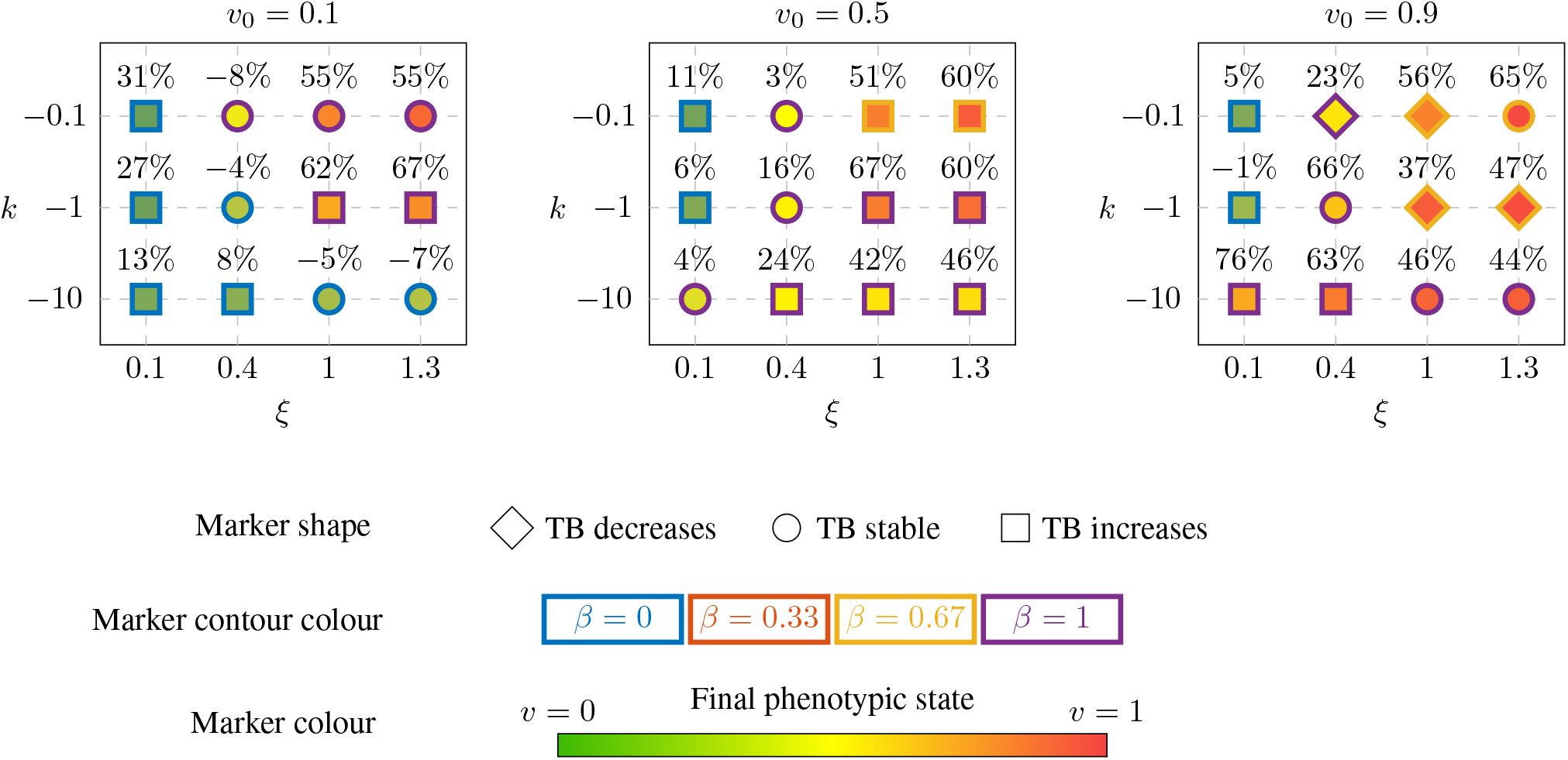
Summary of simulations varying the parameters related with the phenotypic state evolution. Each diagram contains a grid of different shape-location parameters in function *ω* (modelling the acquisition-repair of epigenetic changes that lead to phenotypic plasticity), for a different initial preconditioning condition (that it, a different value of the initial phenotypic state *v*_0_). The marker’s shape determines the global trend of the number of cells (total tumour burden) in each simulation, that is, if the TB tends to increase, decrease of remains stable. The marker’s contour colour represents which value of the heritage parameter *β* favours the most tumour growth and, finally, the marker’s inner colour shows the mean phenotypic state of the tumour cells at the end of the simulation. The percentage above each marker corresponds to the change in number of alive tumour cells during the simulation, when compared with the initial number.

From Figure 3, it can been seen that different values of *k* and *ξ* lead to different global tumour dynamics, due to the nonlinear nature of the model. That is, the impact that the environment has on the phenotypic state of the tumour cells is determinant for the evolution of the tissue. In general, tumours with high *ξ* and *k* (in absolute value) evolve towards CSC phenotypes. Also, a low value of *ξ* generally translates into tumours made of differentiated cells. This is further explored in Figures 4, 5. Besides, lower initial preconditioning (*v*_0_ = 0.1) facilitates the decrease in the number of alive cells.

**Figure 4:**
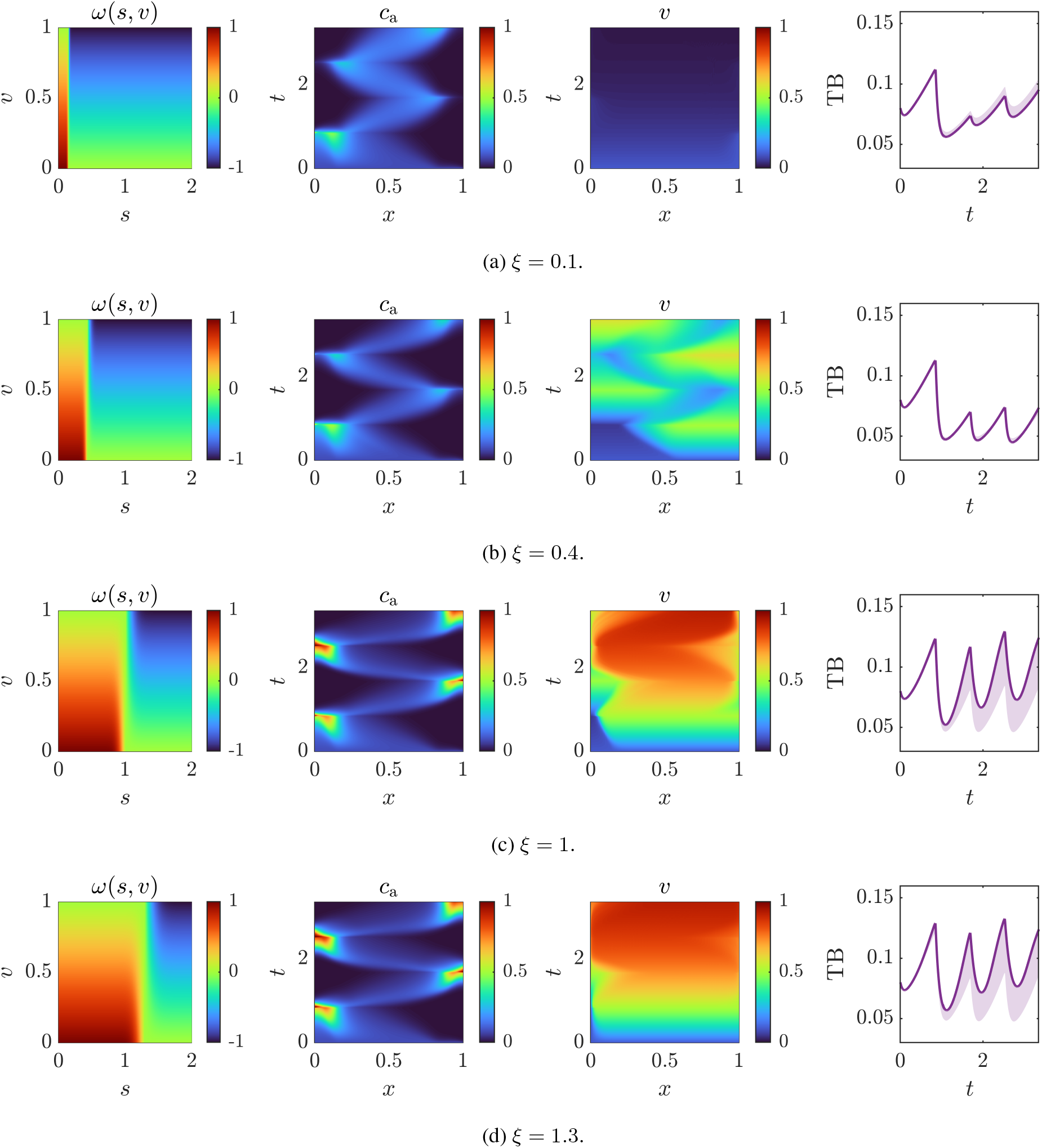
Effect of varying the location parameter *ξ* in function *ω* (*k* = −0.1). The first column represents *ω* as a function of oxygen *s* and the phenotypic state *v*. The second and third columns show the spatio-temporal dynamics of alive cells and the phenotypic state respectively. The fourth column shows the evolution of tumour burden with respect to time, taking into account the variability induced by the heritage parameter *β* (coloured region), with the solid purple line corresponding to the tumour burden in the complete heritage case *β* = 1.

**Figure 5:**
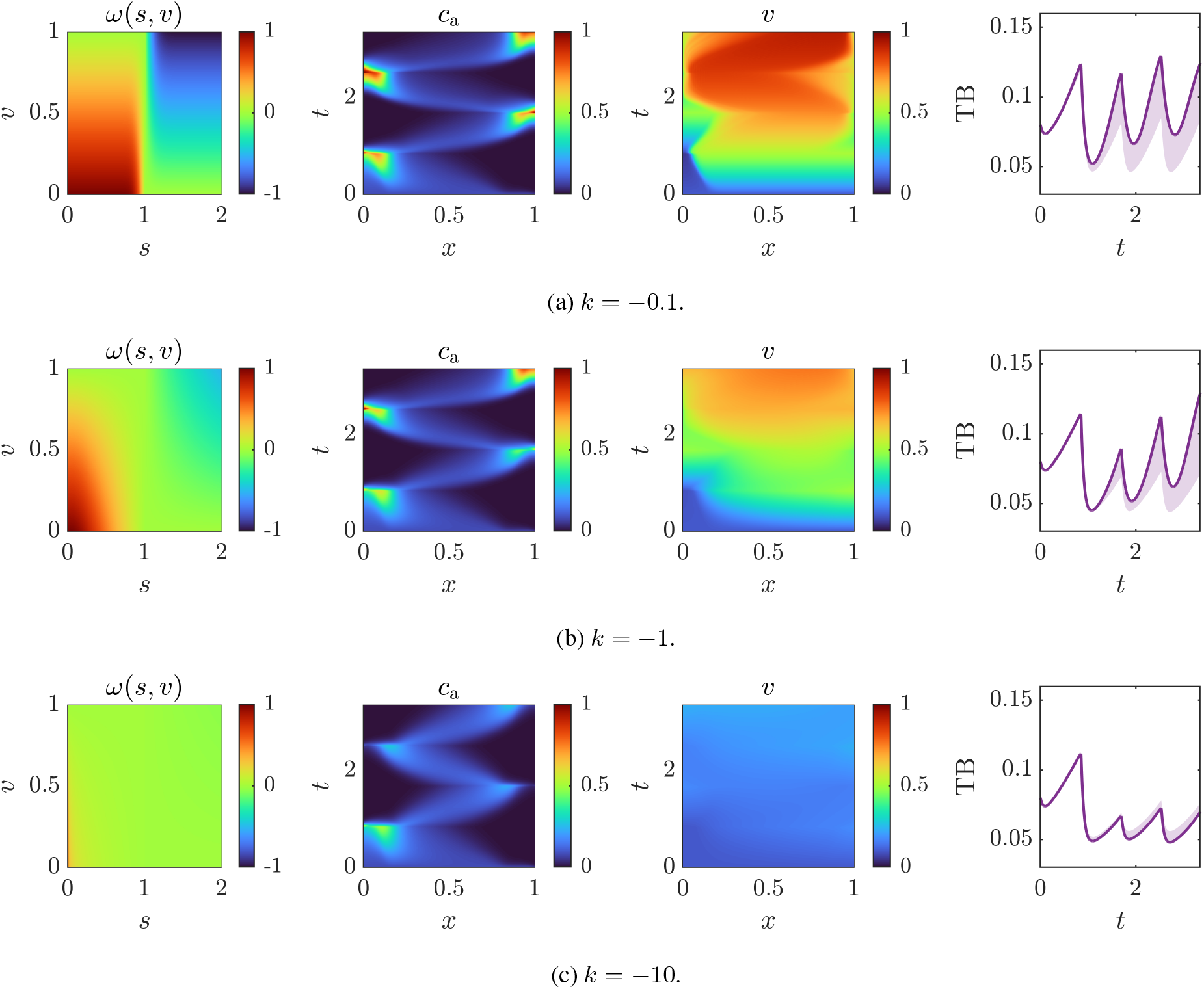
Effect of varying the shape parameter *k* in function *ω* (*ξ* = 1). The first column of figures represents *ω* as a function of oxygen *s* and the phenotypic state *v*. The second and third columns show the spatio-temporal dynamics of alive cells and the phenotypic state respectively. The fourth column shows the evolution of the tumour burden with respect to time, taking into account the variability induced by the heritage parameter *β* (coloured region), with the solid purple line corresponding to the tumour burden in the complete heritage case *β* = 1.

Figure 4 represents the simulation results for different values of *ξ*, leaving *k* = −0.1, *β* = 1 and *v*_0_ = 0.1. *ξ* represents, somehow, the threshold determining if the cell’s phenotype evolve towards a CSC phenotype (*s < ξ*) or towards a differentiated one (*s > ξ*). The higher the parameter *ξ*, the closer to CSC phenotype is the final phenotypic state. This is because a high value of *ξ* implies that cells are driven towards the CSC phenotype for a wider range of oxygen concentrations, that is, they are more sensitive to hypoxia.

Analogously, in Figure 5 we can see simulations corresponding to different values of *k*, leaving *ξ* = 1, *β* = 1 and *v*_0_ = 0.1. This parameter affects the spread of *ω* in the sense that higher absolute values of *k* lead to a smoother *ω* function, where cells are less sensitive to changes in the oxygen concentration. In particular, for *k* = −10 the cells’ phenotype is almost insensitive to the oxygen concentration, so the phenotypic state is only slightly modified throughout the simulation. Indeed, in Figure 3, we can observe that, in all the cases with *k* = −10, the final phenotypic state is similar to the initial one. For lower absolute values of *k*, the tumour is more aggressive in terms of velocities of proliferation and migration.

Phenotypic state heritage also modifies the way the tumour evolves. Assuming *β* = 0 implies that cells, in some sense, repair the epigenetic changes caused by hypoxia through proliferation. At the other end, *β* = 1 implies that the population mean phenotypic state is preserved regardless of cell proliferation. From Figure 3 it can be deduced that when the tumour is composed of differentiated cells (low *v*, greenish marker colours), tumour growth is favoured by no state heritage (*β* = 0). Conversely, for tumours with a phenotypic state that is close to CSC, tumours with *β* = 1 grow more. There are even some cases where an intermediate state conservation by heritage (*β* = 0.67) yields the tumours with the highest number of alive cells, which we can relate with a higher size of the active tumour. This can be explained by the hypotheses regarding the effect of the phenotypic state on cells via the *ψ* functions (see Figure 1). For low values of *v* (phenotype of differentiated cells), overall tumour survival and growth is maximal for *v* = 0, since both death and uptake reach a minimum, compared with low *v* ≠ 0, where death is at its peak. Thus, *β* = 0 helps achieving the *v* = 0 phenotype since, as commented above, it acts as a repair process. Likewise, in the regime of CSCs (high values of *v*), the scenario promoting tumour growth coincides with the peak in *ψ*_gr_ (*v* ≈ 0.8) and *β* = 1 helps cells achieving that range of phenotypic states. Since *ψ*_gr_ is not monotonic, there may be cases where a value of *β <* 1 helps getting the phenotype which best promotes tumour growth.

In Figure 6 we analyse the effect of *β* for two different scenarios (I and II), corresponding to different behaviours, and also for different initial phenotypic states. Scenario I (Figure 6a) corresponds to a case where cells barely change their phenotype. On the other hand, scenario II (Figure 6b) corresponds to a case in which cells’ phenotype is highly sensitive to changes in the oxygen concentration. Consequently, in the former, the phenotypic state mainly changes due to heritage when cells proliferate while, in the latter, cells mainly evolve towards a CSC phenotype throughout successive cycles. In scenario I, with *v*_0_ = 0.1, it can be observed that the tumour is within the differentiated cells regime, and therefore, as commented, *β* = 0 yields the biggest TB. In the rest of cases, cells have phenotypes closer to CSC and the value of *β* that produces the largest tumour is that yielding the closest value to *v* ≃ 0.8, that is, as pointed out before, the value that promotes tumour growth. However, in most cases the value of *β* does not modify the general progression pattern. The only exception is scenario I with *v*_0_ = 0.9, where there are two differentiated trends, *β* = 1 produces a growing tumour with with predominantly CSCs whereas *β <* 0 leads to TB reduction.

**Figure 6:**
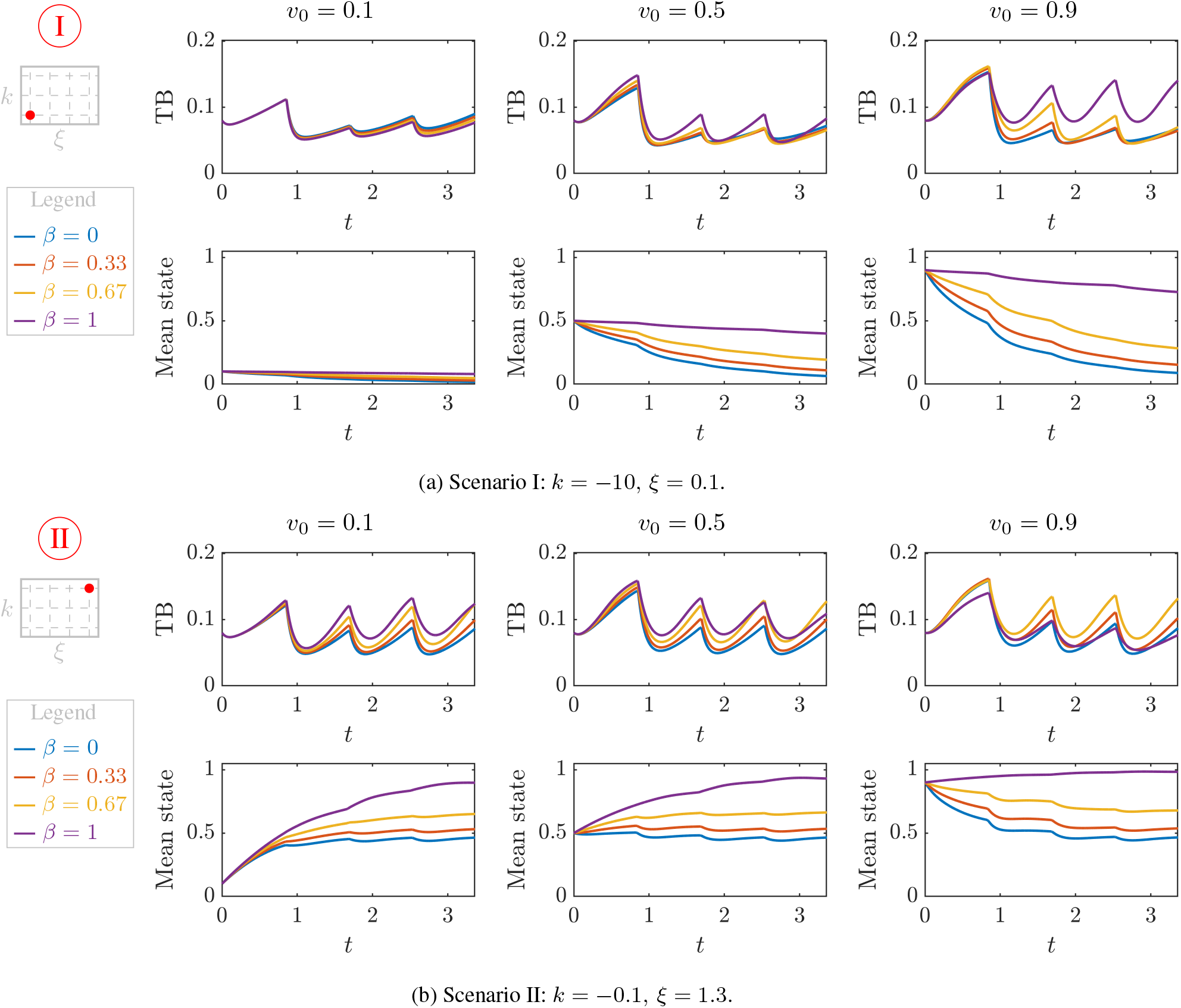
Evolution of the tumour burden (upper row in each subfigure) and the mean phenotypic state (lower row in each subfigure). Simulations consider different values of *β* and *v*_0_, as well as two different combinations of (*k, ξ*) parameters.

Next, we investigate the effect of different initial spatial distributions of the phenotypic state, that is, the effect of having cells distributed in different patterns according to their phenotype. In particular, we analyse four different distributions, all verifying that their mean state is 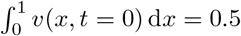:

- uniform *v*_0_ = 0.5,
- left-skewed: step-like distribution, with CSCs in the left (*v*_0_ = 0.9) and differentiated cells in the right (*v*_0_ = 0.1):

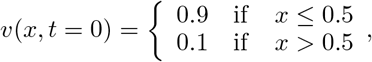
- right-skewed: step-like distribution, with differentiated cells in the left (*v*_0_ = 0.1) and CSCs in the right (*v*_0_ = 0.9):

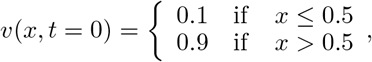
- symmetric and non-uniform: logit-normal distribution with *μ* = 0, *σ* = 0.8:

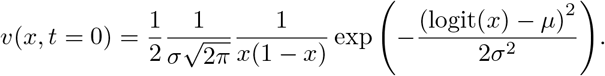

Figures 7 and 8 show the results of TB and mean phenotypic state in two scenarios, for *β* = 0 and *β* = 1 respectively. In the first scenario, in both figures (Figures 7a, 8a), cells can withstand low levels of oxygen without acquiring stemness (the location parameter *ξ* is low), while the opposite occurs in scenario II (Figures 7b, 8b). Overall, it can be seen that although the spatial distribution of the phenotypic state is different in the early simulation times, this does not make a difference in the TB evolution or in the cell spatial distribution, neither qualitatively nor quantitatively. Only in Figure 8a we can appreciate some differences, as an initial *step right* distribution yields a smaller tumour. Even in this case, differences are not significant, so we may conclude that the initial distribution of the phenotypic state does not affect tumour evolution in the long term.

**Figure 7:**
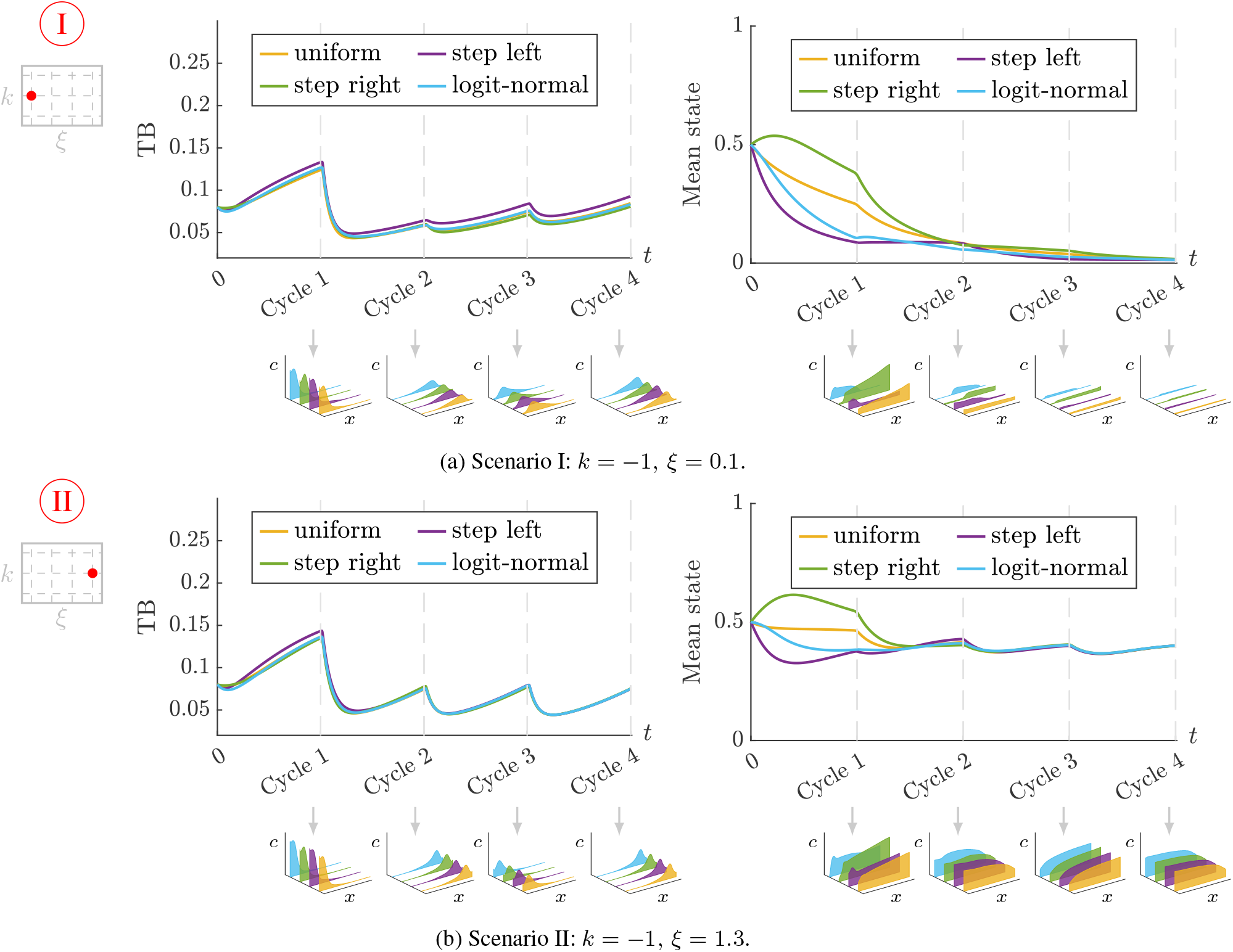
Effect of the initial distribution of cells according to their phenotype on tumour evolution, for the two different scenarios and no inheritance. Scenario I corresponds to cell phenotype insensitive to oxygen variations (*k* = −1, *ξ* = 0.1), and scenario II to cell phenotype insensitive to oxygen variations (*k* = −1, *ξ* = 1.3). The TB and mean state are represented for each scenario, together with the spatial distributions of *c*_a_ and *v* at the end of each cycle.

**Figure 8:**
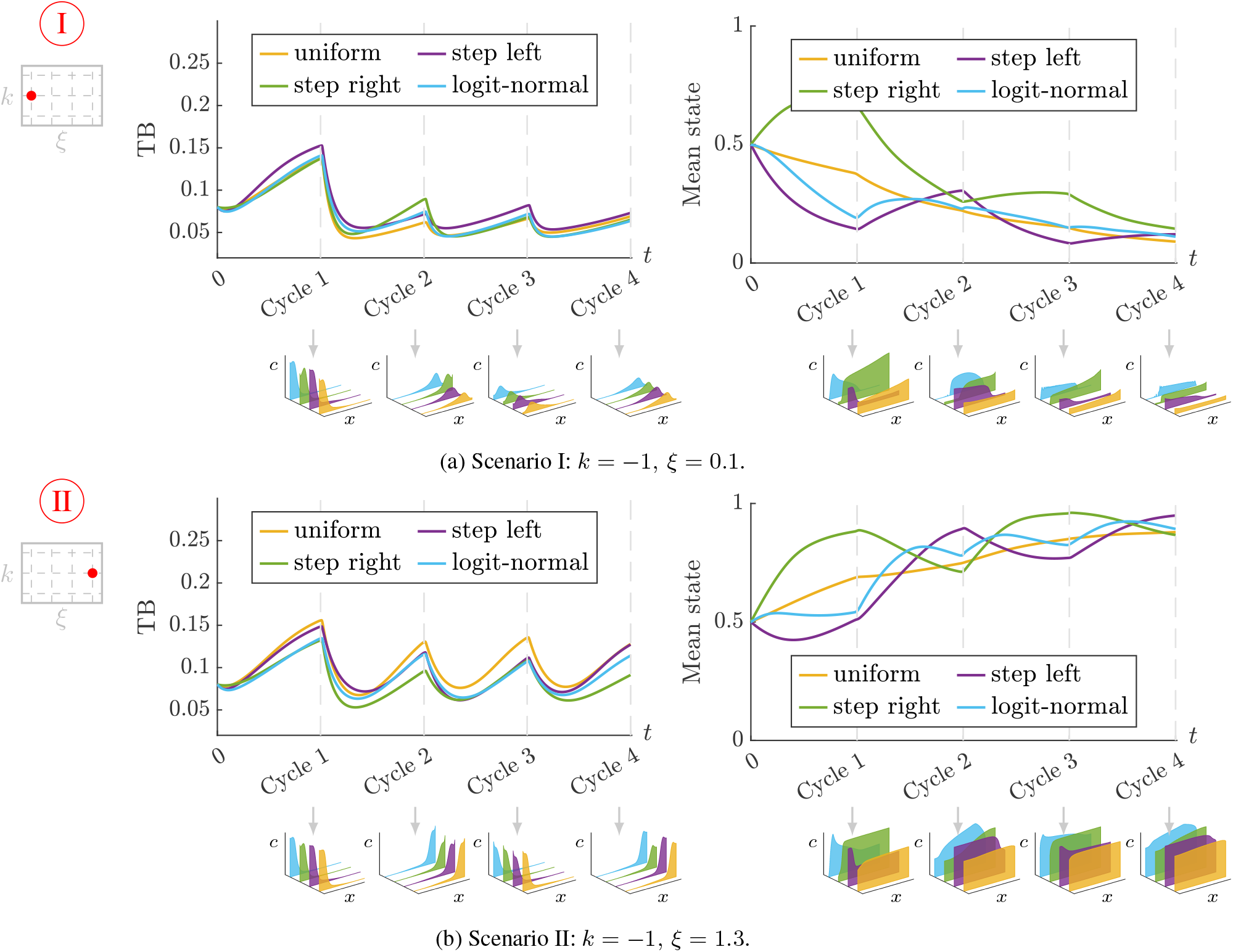
Effect of the initial distribution of cells according to their phenotype on tumour evolution, for the two different scenarios and complete inheritance. Scenario I corresponds to cell phenotype insensitive to oxygen variations (*k* = −1, *ξ* = 0.1), and scenario II to cell phenotype insensitive to oxygen variations (*k* = −1, *ξ* = 1.3). The TB and mean state are represented for each scenario, together with the spatial distributions of *c*_a_ and *v* at the end of each cycle.

To finish this section, we study the effect of modifying the hypotheses on the effect of the phenotypic state in cell behaviour, i.e., the definition of the *ψ* functions in the model. We compare the initial hypotheses defined in Section 2 (Figure 1), which we consider the “base” condition, with a case in which hypoxia only modifies the proliferation capacity of cells, a scenario in which hypoxia modifies everything but the proliferation capacity, and a scenario in which it modifies everything but the oxygen uptake. A summary of these scenarios is presented in Table 2. The simulation results are presented in Figure 9. From this figure, it can be seen that the effect of the phenotypic state on growth is essential to explain the different trends observed, since the assumption that there is no effect on growth leads to tumours that do not grow significantly. Indeed, the increase in growth is one of the defining features of CSCs [24]. On the other hand, considering that the change in phenotype is restricted only to the growth capability always yields bigger tumours. This is not a good representation of reality, since it would lead always to growing tumours (for example, in Figure 9a, it can be seen that the base hypothesis may lead to decreasing tumours, while considering only the effect on growth does not produce that trend). Besides, we know that the effect on motility (via the EMT) is also needed to characterise the transition towards CSCs. Finally, other authors have neglected the effect of phenotypic plasticity on oxygen uptake [36]. As can be seen in Figure 9, this also leads invariably to big growing tumours. This hypothesis is not physiologically consistent as it fails to fulfill energetic constraints. That is, since cells have limited resources, an increase in their metabolic activity, either proliferative or migratory, must lead to an increase in uptake to fulfill the cell energy demands.

**Table 2:** Summary of the different hypotheses regarding the effect of the internal variable in cell behaviour. The different hypotheses are translated into the activation/deactivation of the corresponding *ψ* function, a value of 1 means that the phenotypic state has no effect on that particular phenomenon.

| Scenario | Growth | Motility | Death | Uptake |
| --- | --- | --- | --- | --- |
| Base | $\psi_{gr}$ | $\psi_{mot}$ | $\psi_d$ | $\psi_u$ |
| Effect only on growth | $\psi_{gr}$ | 1 | 1 | 1 |
| No effect on growth | 1 | $\psi_{mot}$ | $\psi_d$ | $\psi_u$ |
| No effect on uptake | $\psi_{gr}$ | $\psi_{mot}$ | $\psi_d$ | 1 |

**Figure 9:**
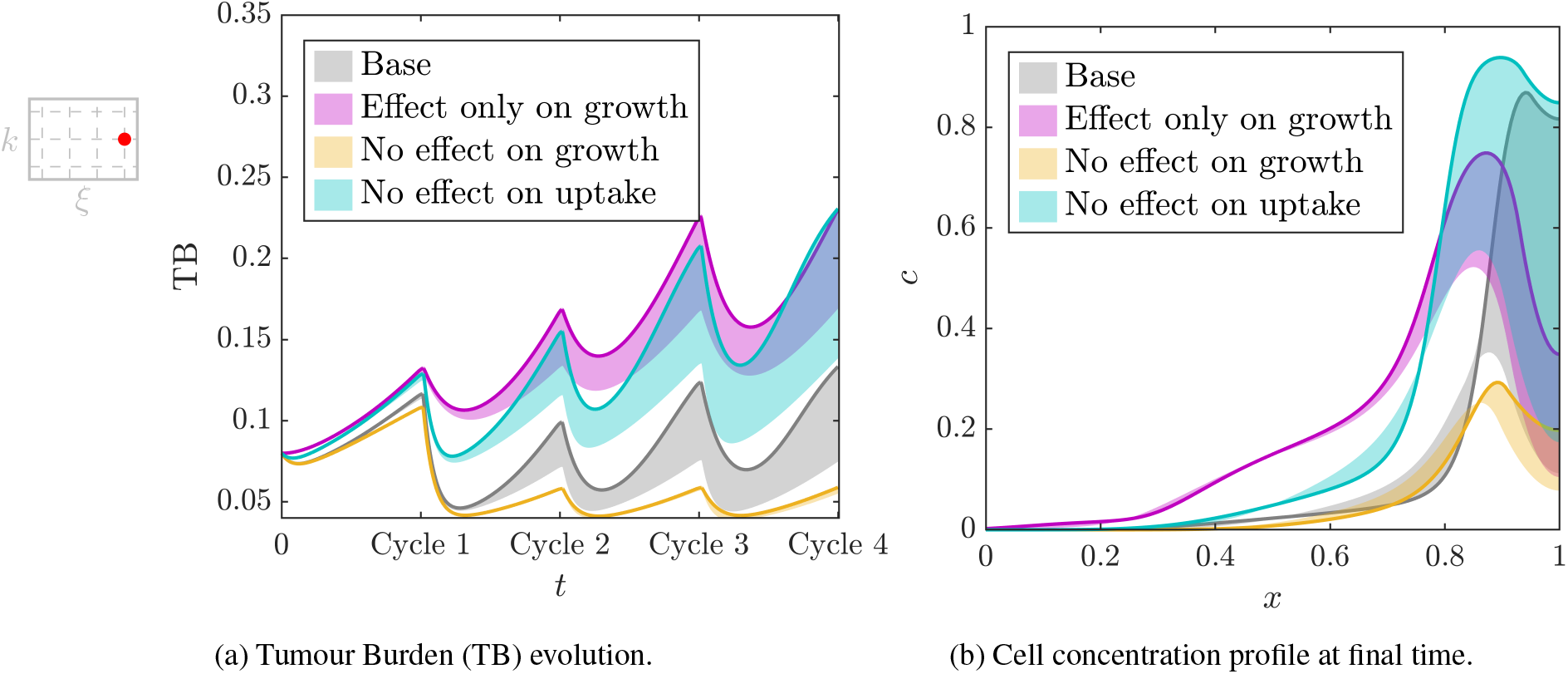
Effect of different hypotheses about the effect of the internal variable in cell behaviour. The results are presented for a moderately sensitive tumour (*k* = −1, *ξ* = 1.3) initially composed of differentiated cells (*v*_0_ = 0.1). The coloured band represents tumour evolution taking into account all possible *β* values, with the solid line corresponding to the simulation with *β* = 1.

## 4 Study of GBM dynamics

In this section, we simulate GBM evolution under different oxygenation conditions. To simulate cyclic hypoxia, we define the (non-dimensional) oxygen concentration at the left (*s*_*L*_) and right (*s*_*R*_) edges of the domain with periodic boundary conditions in the form of sinusoidal functions, that can be parameterised by their period (*𝒯*), minimum level of oxygen (*s*_min_), amplitude (*A*) and phase-shift between both functions (*ϕ*):

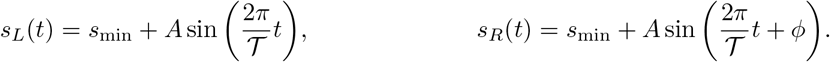

We perform a sensitivity analysis to elucidate the effect of each of these four parameters on tumour evolution. We used Latin Hypercube Sampling (LHS) to generate 300 combinations of the aforementioned parameters, with 0.12 ≤ *𝒯* ≤ 1.4, 0 ≤ *s*_min_ ≤1, 0.14 ≤ *A* ≤ 1.14, 0 ≤ *ϕ* ≤ *π*. Then, for each parameter combination, we simulate the tumour evolution for our two reference scenarios up to *t* = 10 and compute the Partial Correlation Coefficients (PCCs) of each parameter with the TB at the end of the simulation. As observed in Figure 10, in both scenarios, the period has no relevant correlation with the output value (final TB), while the TB has the highest correlation with the minimum value of oxygen *s*_min_.

**Figure 10:**
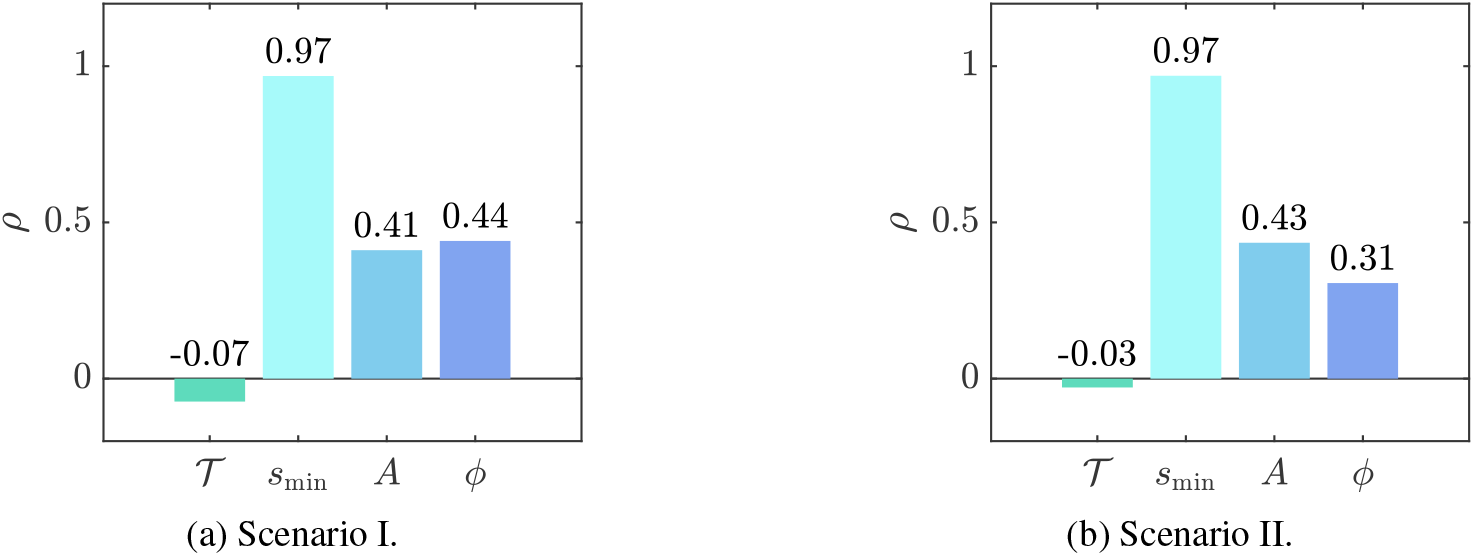
Partial correlation coefficients relating the parameters defining oxygen boundary conditions to the final tumour burden. Scenario I corresponds to cells whose phenotype is quite insensitive to oxygen variation, while in scenario II, the cells’ phenotype is sensitive to changes in oxygen concentration.

In Figure 11 we represent in a scatter plot the different simulations organised in terms of the values of *s*_min_, *A* and *ϕ*. The dot size is proportional to the period *𝒯* (the bigger the dot, the higher the period in that simulation). As previously shown, *𝒯* does not have a significant impact in the value of the final TB, so there is no particular pattern in the distribution of dot sizes in Figure 11. We distinguish three global trends for the tumour burden: i) the TB at the end is greater than at the beginning of the simulation (represented with a blue dot); ii) the TB at the end is smaller than at the beginning of the simulation (represented with a yellow dot); iii) the tumour has remitted at some point in the simulation (represented with an orange dot).

**Figure 11:**
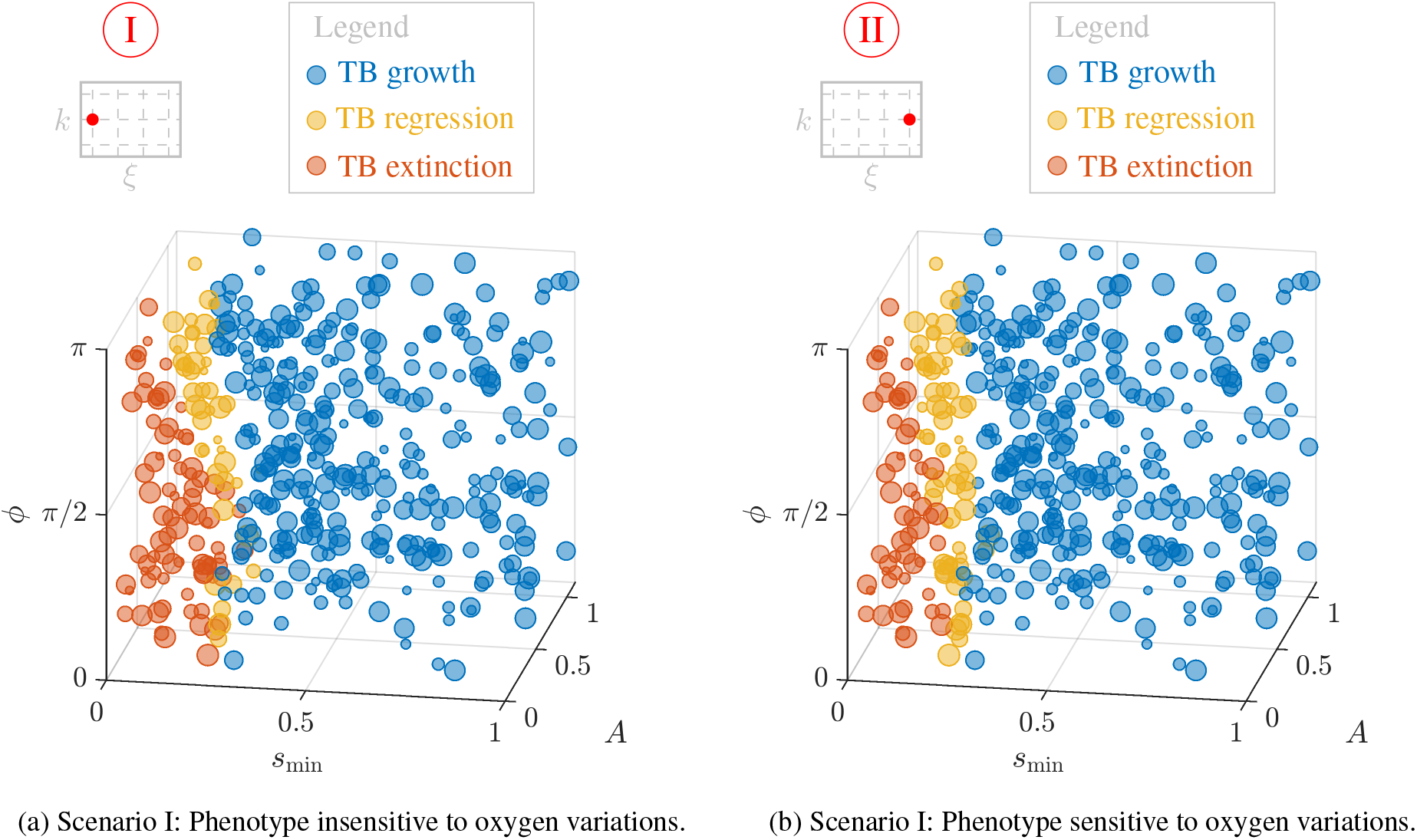
Scatter plot of the outcome of the simulations in terms of *s*_min_, *A* and *ϕ*. A blue dot indicates that the TB has grown, a yellow dot indicates that it has diminished, and an orange dot indicates that the tumour has become extinct. The dot size is proportional to the period *𝒯* so that bigger dots correspond to simulations with a higher period.

It can be seen that for both scenarios the results are similar. As shown by the PCCs, the parameter with the highest influence on TB dynamics is *s*_min_. Besides, lower values of the phase-shift increment the possibilities of tumour remission. In the limits, when *ϕ* → 0 there are times when the whole chamber is under hypoxic conditions, while when *ϕ* → *π* there is always an oxygen gradient, with the more oxygenated area offering higher surviving possibilities.

Interestingly, insensitive tumours show a rate of extinction of 14%, while in the case sensitive this rate drops to 9%, suggesting that our model is able to capture the increased resilience that adaptation confers. To further study this, we calculate the surface defining the ranges of the parameters *s*_min_, *A* and *ϕ* that cause tumour remission. We use Support Vector Machines (SVMs) with a polynomial kernel of order 2, since they provide an easy an efficient tool for nonlinear classifications [66]. The resulting surfaces are represented in Figure 12, where it can be observed that the surface corresponding to insensitive tumours encloses a bigger area of remission. In particular, sensitive tumours seem to be able to withstand more severe hypoxia levels (lower values of *s*_min_), especially for higher amplitudes.

**Figure 12:**
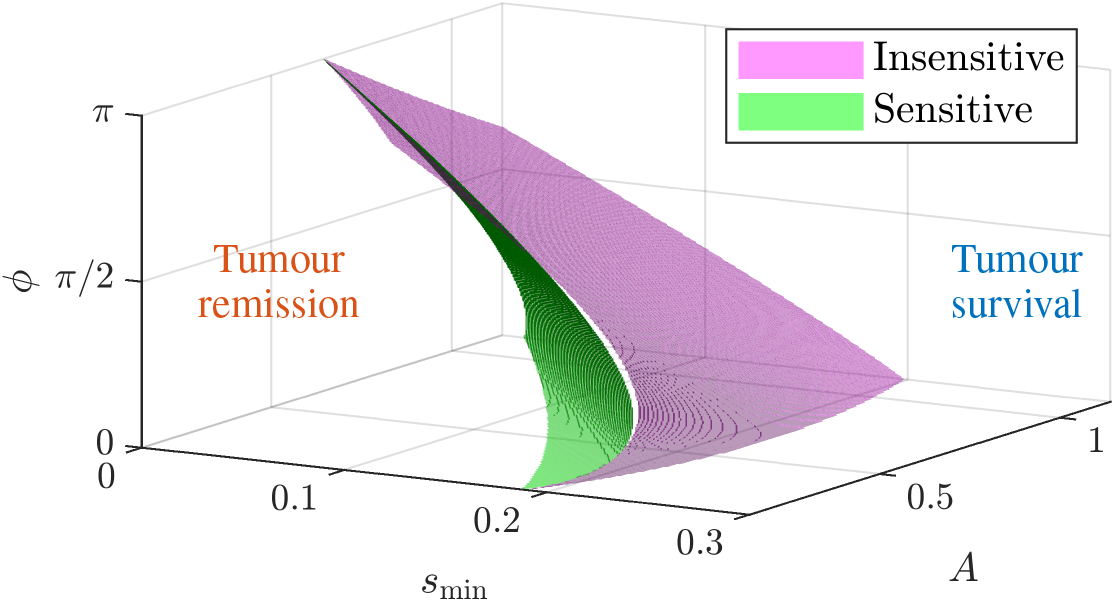
Surface separating tumour remission and tumour survival. This surface has been calculated for sensitive (green) and insensitive (pink) populations using SVM with a polynomial kernel of order 2.

Additional results showing the complete temporal evolution of the TB for some relevant parameter combinations can be found in Appendix **??** (Figure **??**). TB evolution shows that, in general, tumours that do not adapt in response to hypoxia (insensitive tumours) have a greater number of cells, i.e. a higher TB, than the ones that adapt. However, these latter survive longer, showing that even if the adaptive changes do not result in a higher TB, they provide an advantage for surviving in adverse environments.

## 5 Discussion

Throughout this paper, we have presented a new approach to modelling cellular adaptation and phenotypic plasticity in tumour evolution. This approach is based on incorporating internal variables to a previously defined continuum model. Here, internal variables fully describe cell state and its evolution representing, in a macroscopic sense, the molecular pathways that lead to changes in the cell’s phenotype. In turn, cell state controls the response to external stimuli and overall cell behaviour.

In the past years there have been a number of previous models tackling cell adaptation with continuum models from different perspectives. Models with a certain number of discrete phenotypes are the most widespread. In those models, each phenotype has a different behaviour (e.g., different proliferation or migration rate) usually represented by a differential equation, and cells switch from one phenotype to another with a certain transition rate, which is usually mediated by environmental conditions, such as the concentration of oxygen or a drug [35, 37]. These models are simple to implement, because they only imply adding additional equations (one per phenotype), including phenotype transitions and changing the parameters; and they allow to reproduce some important tendencies. However, they do not reproduce the biological reality, where cells do not transition between discrete states, but undergo a range of different behaviours according to their gene expression, as have been shown for different cancers, such as colon [67] or lung [68]. Additionally, the number of parameters increases linearly with the number of different phenotypes considered, thus obscuring the fitting procedure and introducing numerical slack that may lead to overfitting.

There are also some approaches where cell phenotype is modelled using an artificial independent variable *d* ∈ [0, *d*^max^] [36, 34], representing the range of possible cell states, that is, adding an extra dimension to the problem (we will refer to this approach as *external dimension approach*, in opposition to our *internal variable approach*). This method overcomes the aforementioned limitation of discrete-phenotype models. Also, it does not require additional equations, but only the inclusion of flux terms on dimension *d* to account for cell state transitions. Besides, this approach allows obtaining the phenotype spatial distribution at each time point. However, it may be necessary at some point to include different internal variables, due to, for example, different scales for different epigenetic processes, such as the response to hypoxia and to a drug. In this approach, this implies adding extra dimensions, whose interpretation and implementation may be cumbersome, requiring discretisation in higher dimension spaces. This leads to increases in the computational cost of the simulations, giving rise to the so-called curse of dimensionality.

The work by Lei et al. [69] presents another approach to cell adaptation that also takes into account the distribution of phenotypic states at each point, on this occasion for the case of stem cell regeneration. However, the model does not use differential equations but integral equations to evaluate cell number at each time point (there is no spatial coordinate). The model also lacks the relationship between the environment and the phenotype, with cells acquiring epigenetic changes randomly, even if it does consider the inheritance of such states.

The model presented in this paper describes phenotype as a continuum field, but instead of representing it as an extra dimension, we use internal variables to describe it. To the authors’ knowledge, this is the first model in cell evolution taking this approach, which is otherwise widely used in other disciplines, such as damage [44, 70, 71] or control theory [43]. This approach requires defining an evolution equation for the internal variables, which in our case takes the form of another transport equation. State internal variables have been mostly used in Lagrangian frameworks, whereas we introduce them within an Eulerian one. This requires incorporating convective terms, inherent to Eulerian frameworks, to enable cells to keep their state. This feature represents another novelty in the approach of this paper. The definition of the transport equation for cell phenotype is straightforward from the biological hypotheses, allowing to include source terms directly relating stimuli and cell state. This may represent an advantage with respect to the external dimension approach, where interpretability is somehow hindered and of course, with respect to [69], where there is no relationship between state and environment, something crucial for cellular adaptation. In contrast to [69], our model also incorporates the spatial coordinate to account for cell migration, especially relevant for tumour invasion. Finally, sticking to the numerical complexity of the model, including additional internal variables in our model (only) implies adding equations, overcoming the drawbacks related to the computational cost mentioned for the external dimension approach [36]. Also in this regard, it requires less parameters than discrete phenotype models, yielding simpler models, easier to implement, calibrate and validate.

However, the proposed model also presents limitations, since it can only provide the mean phenotypic state at each spatial point, whereas both the external dimension model [36] as well as [69] allow to get the phenotypic distribution at each point. In this sense, our model represents a simplified representation of the models taking acount the phenotypic state distribution [36, 69], and can be seen as a result of averaging the distribution of states at each point. The mathematical relationship between these two different approaches could be further explored in future works.

In summary, the presented framework permits simulating the interplay between cell behaviour and the environment, through the concept of cell state. The model is general and can be adapted to a wide variety of biological problems, with different cell populations, chemical species or any other type of external signal in the surrounding microenvironment and different state or internal variables. The cell state has a unique value at each point. Hence, we loose the state distribution in a local point, but we assume that, at the population level, working with averaged values representing the collective behaviour is a reasonable simplification.

Besides, there are some other limitations related to the validation of the model. As previously said, the model for GBM evolution that is used as a starting point has already been validated with different experiments in microfluidic devices [52, 35, 72] However, the adaptation extension here presented has yet to be experimentally validated. To endow this model with predictive capacity, validating it with experimental data is paramount. Nevertheless, the required experiments are cumbersome and present some important technical difficulties. For example, the cyclic hypoxia experiment in a microfluidic device used here as benchmark requires the ability of opening and closing the channels of the device, and ideally some way of measuring the epigenetic changes that take place, even if qualitatively. Hopefully, microfluidics is a field in constant and rapid evolution, and recently different techniques are emerging to measure epigenetic marks and gene expression via, for example, RNA sequencing [73, 74].

As a first approach to get a higher insight on the performance of the model and to compare with actual biological known facts, we have presented here several illustrative examples with a thorough parametric analysis. The objective of this study is analyzing the potential of the model for capturing important biological trends. Indeed, the results here presented reproduce some trends that have been reported in the existing literature about CSCs, GBM and cyclic hypoxia. It has been reported that the most aggressive tumours contain the highest number of CSCs [59], which is in line with the results shown in Figure 3 where tumours that initially have a higher proportion of CSCs grow more. The model allows obtaining different global trends of tumour evolution depending on the parameters regulating phenotype changes, which reinforces the idea that epigenetic events are behind tumour heterogeneity [26, 75]. Moreover, with this model, different hypotheses regarding the effect of phenotypic changes caused by hypoxia on cell behaviour can be tested, and it has been shown that the effects on growth and oxygen uptake are particularly relevant if we want to reproduce GBM aggressiveness (see Figure 9). This conclusion is in agreement with previous findings, proving that increased growth [60] and consumption [76] are defining features of these cells. Besides, investigating the effect of cyclic hypoxia, we have shown that cells that undergo phenotypic changes to adapt show increased resilience in hypoxic environments (Figures 11, 12). The use of an experiment recreating cyclic hypoxia is supported by the fact that it is a relevant phenomena occurring inside most tumours due to their aberrant vasculature [77], and yielding more aggressive tumours than chronic hypoxia (Figure **??**) [78].

Finally, one-dimensional simulations are carried out as proof of concept of the model, due to its easy implementation and reduced computational cost, but they also imply some limitations. Moreover, the proposed *in silico* experiment, consisting on back and forth migration within the chamber, does not correspond exactly with what happens in the brain, where the tumour has no space constraint and expands. Therefore, the application of this model to real patient-specific geometries is an interesting line to explore in the future.

In spite of these limitations, mathematical models, and in particular the one presented here to model epigenetic changes in cell populations as well as the associated effect on the behaviour of the population, are really useful for understanding biological phenomena. Our model is able to reproduce different trends reported in literature. Of course there is room for improvement, and the validation of the model together with the generalisation to more dimensions and internal variables should be addressed in the future. For example, the inclusion of an internal variable taking into account the phenotypic changes that take place in response to chemotherapy treatments could be interesting, given that there are studies suggesting that drug resistance is favoured by hypoxia [79].

## 6 Conclusion

Cellular adaptation mediated by epigenetic changes and leading to phenotypic heterogeneity is a major concern in cancer research since it has proven to be determinant in the tumour aggressiveness, and hence, in its prognosis, to the extent that it has been included as a Cancer Hallmark in 2022.

In this paper, we have presented a novel approach for modelling these phenomena based on reaction-diffusion equations with internal variables modelling the cell state. This model has been formulated in general, and then particularised to the case of GBM adaptation to hypoxia. Hypoxia promotes the development of CSCs which have increased proliferation and migration rates and are responsible for the dismal prognosis of this tumour.

The main features of the model, such as the acquisition of phenotypic changes, its inheritance and the effect of plasticity in cell behaviour have been analysed by means of an extensive parametric analysis showing the model’s flexibility and capability to get relevant physiological results that capture the huge variability present in tumours. Indeed, it is able to reproduce growing, stable or decreasing tumours, depending on the sensitivity of the cells to changes in their microenvironment and without particular trends, showing the non linearity of the model and the hidden dependencies. We emphasise the importance of taking into account the effect of adaptation to hypoxia in proliferation, because otherwise we are not able to capture the increased aggressiveness observed in literature in tumours subjected to cyclic hypoxia. Besides, the increased consumption caused by phenotypic changes leading to CSCs is also important not to overestimate growth of tumours, neglecting the energetic constraints.

Finally, a study on tumour dynamics, when varying the boundary conditions for oxygen concentrations has been carried out, showing that incorporating cellular adaptation results in an increased resilience of GBM tumours. Indeed, tumours that undergo phenotypic changes are less likely to remit.

The presented framework takes a further step towards creating models that are able to reproduce the complexity of this disease and provide predictive tools to *in silico* test different treatments and conditions.

## Author Contributions

**Marina Pérez-Aliacar:** Conceptualization, Methodology, Software, Formal analysis, Visualization, Writing - Original Draft, Writing - Review & Editing; **Jacobo Ayensa-Jiménez:** Conceptualization, Methodology, Software, Formal analysis, Writing - Review & Editing, Supervision; **Manuel Doblaré:** Conceptualization, Methodology, Writing - Review & Editing, Supervision, Project administration, Funding acquisition;

## Acknowledgements

The authors gratefully acknowledge the financial support from the Spanish Ministry of Science and Innovation (MICINN), the State Research Agency (AEI), and FEDER, UE through the project PID2021-126051OB-C41.

## References

[1] Rebecca J Fox, Jennifer M Donelson, Celia Schunter, Timothy Ravasi, and Juan D Gaitán-Espitia. Beyond buying time: the role of plasticity in phenotypic adaptation to rapid environmental change, 2019.

[2] H Frederik Nijhout. Development and evolution of adaptive polyphenisms. Evolution & development, 5(1):9–18, 2003.

[3] BingKan Xue and Stanislas Leibler. Benefits of phenotypic plasticity for population growth in varying environments. Proceedings of the National Academy of Sciences, 115(50):12745–12750, 2018.

[4] Ben D MacArthur, Avi Ma’ayan, and Ihor R Lemischka. Systems biology of stem cell fate and cellular reprogramming. Nature reviews Molecular cell biology, 10(10):672–681, 2009.

[5] Xinxiu Xu, Quan Wang, Yuan Long, Ru Zhang, Xiaoyuan Wei, Mingzhe Xing, Haifeng Gu, and Xin Xie. Stress-mediated p38 activation promotes somatic cell reprogramming. Cell research, 23(1):131–141, 2013.

[6] JK Kim, Mala Samaranayake, and Sriharsa Pradhan. Epigenetic mechanisms in mammals. Cellular and molecular life sciences, 66:596–612, 2009.

[7] Elizabeth J Duncan, Peter D Gluckman, and Peter K Dearden. Epigenetics, plasticity, and evolution: How do we link epigenetic change to phenotype? Journal of Experimental Zoology Part B: Molecular and Developmental Evolution, 322(4):208–220, 2014.

[8] Sanket Shah, Mudasir Rashid, Tripti Verma, and Sanjay Gupta. Chromatin, histones, and histone modifications in health and disease. Genome Plasticity in Health and Disease, pages 109–135, 2020.

[9] Fabio Mohn and Dirk Schübeler. Genetics and epigenetics: stability and plasticity during cellular differentiation. Trends in Genetics, 25(3):129–136, 2009.

[10] Riya R Kanherkar, Naina Bhatia-Dey, and Antonei B Csoka. Epigenetics across the human lifespan. Frontiers in cell and developmental biology, 2:49, 2014.

[11] Andrew P Feinberg. Phenotypic plasticity and the epigenetics of human disease. Nature, 447(7143):433–440, 2007.

[12] Liping Chen, Weifang Huang, Ling Wang, Zili Zhang, Feng Zhang, Shizhong Zheng, and Desong Kong. The effects of epigenetic modification on the occurrence and progression of liver diseases and the involved mechanism. Expert Review of Gastroenterology & Hepatology, 14(4):259–270, 2020.

[13] Peter A Jones and Stephen B Baylin. The fundamental role of epigenetic events in cancer. Nature reviews genetics, 3(6):415–428, 2002.

[14] Stephen B Baylin and Peter A Jones. A decade of exploring the cancer epigenome—biological and translational implications. Nature Reviews Cancer, 11(10):726–734, 2011.

[15] Antara Biswas and Subhajyoti De. Drivers of dynamic intratumor heterogeneity and phenotypic plasticity. American Journal of Physiology-Cell Physiology, 320(5):C750–C760, 2021.

[16] Douglas Hanahan. Hallmarks of cancer: new dimensions. Cancer discovery, 12(1):31–46, 2022.

[17] Piyush B Gupta, Ievgenia Pastushenko, Adam Skibinski, Cedric Blanpain, and Charlotte Kuperwasser. Phenotypic plasticity: driver of cancer initiation, progression, and therapy resistance. Cell stem cell, 24(1):65–78, 2019.

[18] Thomas Brabletz, Raghu Kalluri M Angela Nieto, and Robert A Weinberg. Emt in cancer. Nature Reviews Cancer, 18(2):128–134, 2018.

[19] Sugandha Bhatia, Peiyu Wang, Alan Toh, and Erik W Thompson. New insights into the role of phenotypic plasticity and emt in driving cancer progression. Frontiers in molecular biosciences, 7:71, 2020.

[20] Hans Clevers. The cancer stem cell: premises, promises and challenges. Nature medicine, 17(3):313–319, 2011.

[21] Lia Walcher, Ann-Kathrin Kistenmacher, Huizhen Suo, Reni Kitte, Sarah Dluczek, Alexander Strauß, André-René Blaudszun, Tetyana Yevsa, Stephan Fricke, and Uta Kossatz-Boehlert. Cancer stem cells—origins and biomarkers: perspectives for targeted personalized therapies. Frontiers in immunology, 11:1280, 2020.

[22] Jeremy N Rich. Cancer stem cells: understanding tumor hierarchy and heterogeneity. Medicine, 95(Suppl 1), 2016.

[23] Christina Scheel and Robert A Weinberg. Phenotypic plasticity and epithelial-mesenchymal transitions in cancer and normal stem cells? International journal of cancer, 129(10):2310–2314, 2011.

[24] Ain Zubaidah Ayob and Thamil Selvee Ramasamy. Cancer stem cells as key drivers of tumour progression. Journal of biomedical science, 25:1–18, 2018.

[25] Richard P Hill, Delphine T Marie-Egyptienne, and David W Hedley. Cancer stem cells, hypoxia and metastasis. In Seminars in radiation oncology, volume 19, pages 106–111. Elsevier, 2009.

[26] Håkan Axelson, Erik Fredlund, Marie Ovenberger, Göran Landberg, and Sven Påhlman. Hypoxia-induced dedifferentiation of tumor cells–a mechanism behind heterogeneity and aggressiveness of solid tumors. In Seminars in cell & developmental biology, volume 16, pages 554–563. Elsevier, 2005.

[27] JM Heddleston, Z Li, JDet al Lathia, S Bao, AB Hjelmeland, and JN Rich. Hypoxia inducible factors in cancer stem cells. British journal of cancer, 102(5):789–795, 2010.

[28] Tijana Stanković, Teodora Randelović, Miodrag Dragoj, Sonja Stojković Burić Luis Fernández, Ignacio Ochoa, Victor M Pérez-García, and Milica Pešić. In vitro biomimetic models for glioblastoma-a promising tool for drug response studies. Drug Resistance Updates, 55:100753, 2021.

[29] Ana Rita Monteiro, Richard Hill, Geoffrey J Pilkington, and Patrícia A Madureira. The role of hypoxia in glioblastoma invasion. Cells, 6(4):45, 2017.

[30] Daniel Bergman and Trachette L Jackson. Phenotype switching in a global method for agent-based models of biological tissue. Plos one, 18(2):e0281672, 2023.

[31] Vito Quaranta, Katarzyna A Rejniak, Philip Gerlee, and Alexander RA Anderson. Invasion emerges from cancer cell adaptation to competitive microenvironments: quantitative predictions from multiscale mathematical models. In Seminars in Cancer Biology, volume 18, pages 338–348. Elsevier, 2008.

[32] Robert A Gatenby, Kieran Smallbone, Philip K Maini, Fabrice Rose, J Averill, Raymond B Nagle, L Worrall, and Robert J Gillies. Cellular adaptations to hypoxia and acidosis during somatic evolution of breast cancer. British journal of cancer, 97(5):646–653, 2007.

[33] Aleksandra Ardaševa, Robert A Gatenby, Alexander RA Anderson, Helen M Byrne, Philip K Maini, and Tommaso Lorenzi. A mathematical dissection of the adaptation of cell populations to fluctuating oxygen levels. Bulletin of Mathematical Biology, 82:1–24, 2020.

[34] Arran Hodgkinson, Laurent Le Cam, Dumitru Trucu, and Ovidiu Radulescu. Spatio-genetic and phenotypic modelling elucidates resistance and re-sensitisation to treatment in heterogeneous melanoma. Journal of theoretical biology, 466:84–105, 2019.

[35] Jose M Ayuso, Rosa Monge, Alicia Martínez-González, María Virumbrales-Muñoz, Guillermo A Llamazares, Javier Berganzo, Aurelio Hernández-Laín, Jorge Santolaria, Manuel Doblaré, Christopher Hubert, et al. Glioblastoma on a microfluidic chip: Generating pseudopalisades and enhancing aggressiveness through blood vessel obstruction events. Neuro-oncology, 19(4):503–513, 2017.

[36] Giulia L Celora, Helen M Byrne, and PG Kevrekidis. Spatio-temporal modelling of phenotypic heterogeneity in tumour tissues and its impact on radiotherapy treatment. Journal of Theoretical Biology, 556:111248, 2023.

[37] Sharon S Hori, Ling Tong, Srividya Swaminathan, Mariola Liebersbach, Jingjing Wang, Sanjiv S Gambhir, and Dean W Felsher. A mathematical model of tumor regression and recurrence after therapeutic oncogene inactivation. Scientific reports, 11(1):1–14, 2021.

[38] Edouard Ollier, Pauline Mazzocco, Damien Ricard, Gentian Kaloshi, Ahmed Idbaih, Agusti Alentorn, Dimitri Psimaras, Jérôme Honnorat, Jean-Yves Delattre, Emmanuel Grenier, et al. Analysis of temozolomide resistance in low-grade gliomas using a mechanistic mathematical model. Fundamental & clinical pharmacology, 31(3):347–358, 2017.

[39] James M Greene, Jana L Gevertz, and Eduardo D Sontag. Mathematical approach to differentiate spontaneous and induced evolution to drug resistance during cancer treatment. JCO clinical cancer informatics, 3:1–20, 2019.

[40] Jacobo Ayensa-Jiménez, Mohamed H Doweidar, Teodora Randelovic Luis J Fernández, Sara Oliván, Ignacio Ochoa, and Manuel Doblaré. On the simulation of organ-on-chip cell processes: Application to an in vitro model of glioblastoma evolution. In Advances in Biomechanics and Tissue Regeneration, pages 313–341. Elsevier, 2019.

[41] Jacobo Ayensa Jiménez, Manuel Doblaré Castellano, and Mohamed Doweidar Mohyeldin. Study of the effect of the tumour microenvironment on cell response using a combined simulation and machine learning approach. application to the evolution of glioblastoma. 2022.

[42] Samantha A Morris. The evolving concept of cell identity in the single cell era. Development, 146(12):dev169748, 2019.

[43] William L Brogan. Modern control theory. Pearson education india, 1991.

[44] Mark F Horstemeyer and Douglas J Bammann. Historical review of internal state variable theory for inelasticity. International Journal of Plasticity, 26(9):1310–1334, 2010.

[45] Mingzhou Guo, Yaojun Peng, Aiai Gao, Chen Du, and James G Herman. Epigenetic heterogeneity in cancer. Biomarker research, 7(1):1–19, 2019.

[46] Jessica Wright. Epigenetics: reversible tags. Nature, 498(7455):S10–S11, 2013.

[47] VA Blomen and J Boonstra. Stable transmission of reversible modifications: maintenance of epigenetic information through the cell cycle. Cellular and Molecular Life Sciences, 68(1):27–44, 2011.

[48] Hao Wu and Yi Zhang. Reversing dna methylation: mechanisms, genomics, and biological functions. Cell, 156(1-2):45–68, 2014.

[49] Irene Lacal and Rossella Ventura. Epigenetic inheritance: concepts, mechanisms and perspectives. Frontiers in molecular neuroscience, 11:292, 2018.

[50] Kostas A Triantaphyllopoulos, Ioannis Ikonomopoulos, and Andrew J Bannister. Epigenetics and inheritance of phenotype variation in livestock. Epigenetics & chromatin, 9(1):1–18, 2016.

[51] Dragan Stajic and Lars ET Jansen. Empirical evidence for epigenetic inheritance driving evolutionary adaptation. Philosophical Transactions of the Royal Society B, 376(1826):20200121, 2021.

[52] Jacobo Ayensa-Jiménez, Marina Pérez-Aliacar, Teodora Randelovic, Sara Oliván, Luis Fernández, José Antonio Sanz-Herrera, Ignacio Ochoa, Mohamed H Doweidar, and Manuel Doblaré. Mathematical formulation and parametric analysis of in vitro cell models in microfluidic devices: application to different stages of glioblastoma evolution. Scientific Reports, 10(1):1–21, 2020.

[53] Haralampos Hatzikirou, David Basanta, Matthias Simon, K Schaller, and Andreas Deutsch. ‘go or grow’: the key to the emergence of invasion in tumour progression? Mathematical medicine and biology: a journal of the IMA, 29(1):49–65, 2012.

[54] Brian Stramer and Roberto Mayor. Mechanisms and in vivo functions of contact inhibition of locomotion. Nature reviews Molecular cell biology, 18(1):43–55, 2017.

[55] Aalpen A Patel, Edward T Gawlinski, Susan K Lemieux, and Robert A Gatenby. A cellular automaton model of early tumor growth and invasion: the effects of native tissue vascularity and increased anaerobic tumor metabolism. Journal of theoretical biology, 213(3):315–331, 2001.

[56] John M Heddleston, Zhizhong Li, Roger E McLendon, Anita B Hjelmeland, and Jeremy N Rich. The hypoxic microenvironment maintains glioblastoma stem cells and promotes reprogramming towards a cancer stem cell phenotype. Cell cycle, 8(20):3274–3284, 2009.

[57] Eduard Batlle and Hans Clevers. Cancer stem cells revisited. Nature medicine, 23(10):1124–1134, 2017.

[58] Eli E Bar. Glioblastoma, cancer stem cells and hypoxia. Brain Pathology, 21(2):119–129, 2011.

[59] Luca Persano, Elena Rampazzo, Giuseppe Basso, and Giampietro Viola. Glioblastoma cancer stem cells: role of the microenvironment and therapeutic targeting. Biochemical pharmacology, 85(5):612–622, 2013.

[60] Justin D Lathia, Stephen C Mack, Erin E Mulkearns-Hubert, Claudia LL Valentim, and Jeremy N Rich. Cancer stem cells in glioblastoma. Genes & development, 29(12):1203–1217, 2015.

[61] Barbara Ortensi, Matteo Setti, Daniela Osti, and Giuliana Pelicci. Cancer stem cell contribution to glioblastoma invasiveness. Stem cell research & therapy, 4(1):1–11, 2013.

[62] Gioacchin Iannolo, Concetta Conticello, Lorenzo Memeo, and Ruggero De Maria. Apoptosis in normal and cancer stem cells. Critical reviews in oncology/hematology, 66(1):42–51, 2008.

[63] Alina Filatova, Till Acker, and Boyan K Garvalov. The cancer stem cell niche (s): the crosstalk between glioma stem cells and their microenvironment. Biochimica et Biophysica Acta (BBA)-General Subjects, 1830(2):2496–2508, 2013.

[64] Erina Vlashi, Chann Lagadec, Laurent Vergnes, Tomoo Matsutani, Kenta Masui, Maria Poulou, Ruxandra Popescu, Lorenza Della Donna, Patrick Evers, Carmen Dekmezian, et al. Metabolic state of glioma stem cells and nontumorigenic cells. Proceedings of the National Academy of Sciences, 108(38):16062–16067, 2011.

[65] Robert D Skeel and Martin Berzins. A method for the spatial discretization of parabolic equations in one space variable. SIAM journal on scientific and statistical computing, 11(1):1–32, 1990.

[66] Martin Hofmann. Support vector machines-kernels and the kernel trick. Notes, 26(3):1–16, 2006.

[67] Winston R Becker, Stephanie A Nevins, Derek C Chen, Roxanne Chiu, Aaron M Horning, Tuhin K Guha, Rozelle Laquindanum, Meredith Mills, Hassan Chaib, Uri Ladabaum, et al. Single-cell analyses define a continuum of cell state and composition changes in the malignant transformation of polyps to colorectal cancer. Nature genetics, 54(7):985–995, 2022.

[68] Sarah Maddox Groves, Abbie Ireland, Qi Liu, Alan J Simmons, Ken Lau, Wade T Iams, Darren Tyson, Christine M Lovly, Trudy G Oliver, and Vito Quaranta. Cancer hallmarks define a continuum of plastic cell states between small cell lung cancer archetypes. BioRxiv, pages 2021–01, 2021.

[69] Jinzhi Lei, Simon A Levin, and Qing Nie. Mathematical model of adult stem cell regeneration with cross-talk between genetic and epigenetic regulation. Proceedings of the National Academy of Sciences, 111(10):E880–E887, 2014.

[70] Sumio Murakami. Continuum damage mechanics: a continuum mechanics approach to the analysis of damage and fracture, volume 185. Springer Science & Business Media, 2012.

[71] Gérard A Maugin. The saga of internal variables of state in continuum thermo-mechanics (1893–2013). Mechanics Research Communications, 69:79–86, 2015.

[72] Jose M Ayuso, María Virumbrales-Muñoz, Alodia Lacueva, Pilar M Lanuza, Elisa Checa-Chavarria, Pablo Botella, Eduardo Fernández, Manuel Doblare, Simon J Allison, Roger M Phillips, et al. Development and characterization of a microfluidic model of the tumour microenvironment. Scientific reports, 6(1):1–16, 2016.

[73] Yu Hou, Huahu Guo, Chen Cao, Xianlong Li, Boqiang Hu, Ping Zhu, Xinglong Wu, Lu Wen, Fuchou Tang, Yanyi Huang, et al. Single-cell triple omics sequencing reveals genetic, epigenetic, and transcriptomic heterogeneity in hepatocellular carcinomas. Cell research, 26(3):304–319, 2016.

[74] Boyko Kakaradov, Janilyn Arsenio, Christella E Widjaja, Zhaoren He, Stefan Aigner, Patrick J Metz, Bingfei Yu, Ellen J Wehrens, Justine Lopez, Stephanie H Kim, et al. Early transcriptional and epigenetic regulation of cd8+ t cell differentiation revealed by single-cell rna sequencing. Nature immunology, 18(4):422–432, 2017.

[75] Mark Shackleton, Elsa Quintana, Eric R Fearon, and Sean J Morrison. Heterogeneity in cancer: cancer stem cells versus clonal evolution. Cell, 138(5):822–829, 2009.

[76] Michalina Janiszewska, Mario L Suvà, Nicolo Riggi, Riekelt H Houtkooper, Johan Auwerx, Virginie Clément-Schatlo, Ivan Radovanovic, Esther Rheinbay, Paolo Provero, and Ivan Stamenkovic. Imp2 controls oxidative phosphorylation and is crucial for preserving glioblastoma cancer stem cells. Genes & development, 26(17):1926–1944, 2012.

[77] Carine Michiels, Céline Tellier, and Olivier Feron. Cycling hypoxia: A key feature of the tumor microenvironment. Biochimica et Biophysica Acta (BBA)-Reviews on Cancer, 1866(1):76–86, 2016.

[78] Chia-Hung Hsieh, Woei-Cherng Shyu, Chien-Yi Chiang, Jung-Wen Kuo, Wu-Chung Shen, and Ren-Shyan Liu. Nadph oxidase subunit 4-mediated reactive oxygen species contribute to cycling hypoxia-promoted tumor progression in glioblastoma multiforme. PloS one, 6(9):e23945, 2011.

[79] Chii-Wen Chou, Chi-Chung Wang, Chung-Pu Wu, Yu-Jung Lin, Yu-Chun Lee, Ya-Wen Cheng, and Chia-Hung Hsieh. Tumor cycling hypoxia induces chemoresistance in glioblastoma multiforme by upregulating the expression and function of abcb1. Neuro-oncology, 14(10):1227–1238, 2012.

