## Supplementary material for "Modelling cell adaptation using internal variables: accounting for cell plasticity in continuum mathematical biology"

### A Derivation of the evolution equations.

The main equations of the mathematical models presented throughout this paper are differential conservation equations, which are derived from their integral form.

#### A.1 Cells and chemical species

Let us suppose that we have $m$ cell populations whose concentration is $C_i$, and $n$ chemical species with concentration $S_i$. We may define a vector $\boldsymbol{U}$ formed by all the $m+n$ solution variables as:

$$\boldsymbol{U} = [C_1, \cdots, C_m, S_1, \cdots, S_n], \qquad i = 1, \cdots, n+m. \tag{1}$$

The integral conservation law for any of the variables defined per unit volume may be written as:

$$\frac{\mathrm{d}}{\mathrm{d}t}\int_{\mathcal{V}} U_i \,\mathrm{d}v = -\oint_{\partial\mathcal{V}} \boldsymbol{q}_i \cdot \mathrm{d}\boldsymbol{S} + \int_{\mathcal{V}} F_i \,\mathrm{d}v, \tag{2}$$

where $\boldsymbol{q}_i$ stands for the flux of cells/chemical species that goes through the surface of the control volume $\mathcal{V}$, and $F_i$ represents the source term in each time per unit volume.

Applying the divergence theorem, we get:

$$\frac{\mathrm{d}}{\mathrm{d}t}\int_{\mathcal{V}} U_i \,\mathrm{d}v = -\int_{\mathcal{V}} \boldsymbol{\nabla}\cdot \boldsymbol{q}_i \,\mathrm{d}v + \int_{\mathcal{V}} F_i \,\mathrm{d}v, \tag{3}$$

from where we can obtain, for an arbitrary volume $\mathcal{V}$:

$$\int_{\mathcal{V}} \left( \frac{\partial U_i}{\partial t} + \boldsymbol{\nabla}\cdot \boldsymbol{q}_i - F_i \right) \mathrm{d}v = 0, \tag{4}$$

so finally we derive the differential conservation equation:

$$\frac{\partial U_i}{\partial t} = -\boldsymbol{\nabla}\cdot \boldsymbol{q}_i + F_i, \tag{5}$$

which constitute the system of governing equations in the presented model.

The source term for cells $F_i$ $(i = 1, \cdots, m)$, which will be needed for the derivation of the governing equation for internal variables, comprises the terms of cell proliferation and death, and can be expressed as:

$$F_i = U_i(f_i - d_i). \tag{6}$$

#### A.2 Internal variables

To derive the evolution equation for internal variables, the starting point is the conservation law for the total amount of variable in a Representative Volume Element (RVE).

Let $V_{ik}$ be the $k^{\text{th}}$ internal variable affecting the $i^{\text{th}}$ cell population, whose associated field variable is the cell concentration $U_i$ $(i = 1, \cdots, m)$. Therefore, the total amount of internal variable per unit volume is expressed as $U_i V_{ik}$ and its integral balance equation is:

$$\frac{\mathrm{d}}{\mathrm{d}t}\int_{\mathcal{V}} U_i V_{ik} \,\mathrm{d}v = -\oint_{\partial\mathcal{V}} V_{ik}\boldsymbol{q}_i \cdot \mathrm{d}\boldsymbol{S} + \int_{\mathcal{V}} R_{ik} \,\mathrm{d}v, \qquad i = 1, \cdots, m; \;\; k = 1, \cdots, r_i. \tag{7}$$

where $r_i$ is the number of internal variables affecting the $i^{\text{th}}$ population. The first term of the RHS represents the amount of internal variable going in or out of the volume $\mathcal{V}$ due to the effect of the cell flux $\boldsymbol{q}_i$, and the second term is the source term.

Applying the divergence theorem to the surface integral:

$$\frac{\mathrm{d}}{\mathrm{d}t}\int_{\mathcal{V}} U_i V_{ik} \,\mathrm{d}v = -\int_{\mathcal{V}} \boldsymbol{\nabla}\cdot (V_{ik}\boldsymbol{q}_i) \,\mathrm{d}v + \int_{\mathcal{V}} R_{ik} \,\mathrm{d}v. \tag{8}$$

Letting the temporal derivative inside the integral in the LHS and subsequently grouping the integrals, we obtain:

$$\frac{\partial (U_i V_{ik})}{\partial t} = -\boldsymbol{\nabla}\cdot (V_{ik}\boldsymbol{q}_i) + R_{ik}. \tag{9}$$

Then, applying the product rule:

$$U_i \frac{\partial V_{ik}}{\partial t} + V_{ik} \frac{\partial U_i}{\partial t} = -V_{ik} \boldsymbol{\nabla} \cdot \boldsymbol{q}_i - \boldsymbol{q}_i \cdot \boldsymbol{\nabla} V_{ik} + R_{ik}. \tag{10}$$

and using Eq. (5):

$$U_i \frac{\partial V_{ik}}{\partial t} + V_{ik} \left( -\boldsymbol{\nabla} \cdot \boldsymbol{q}_i + F_i \right) = -V_{ik} \boldsymbol{\nabla} \cdot \boldsymbol{q}_i - \boldsymbol{q}_i \cdot \boldsymbol{\nabla} V_{ik} + R_{ik}. \tag{11}$$

Reorganising and simplifying the previous equation, we arrive to a differential equation for the evolution of the internal variables:

$$U_i \frac{\partial V_{ik}}{\partial t} = -\boldsymbol{q}_i \cdot \boldsymbol{\nabla} V_{ik} + R_{ik} - V_{ik} F_i, \tag{12}$$

where we can introduce the expression for $F_i$ defined in Eq. (6):

$$U_i \frac{\partial V_{ik}}{\partial t} = -\boldsymbol{q}_i \cdot \boldsymbol{\nabla} V_{ik} + R_{ik} - V_{ik} U_i (f_i - d_i), \tag{13}$$

In Eq. (13) it remains to be defined the source term for the internal variables $R_{ik}$, defining the amount of $V_{ik}$ that is generated or disappears at each time moment. $R_{ik}$ is presented as a sum of different terms $R_{ik} = \sum_n R_{ik}^n$, comprising the following phenomena:

- $R_{ik}^1$: Internal variable accumulation due to external effects. One of the main hypothesis of this framework is that the phenotypic state of a cell and its associated behaviour depend on the history of external stimuli to which the cell has been subjected. Therefore, the internal variable representing this state depends on those external stimuli through function $\Omega_{ik}$, which may depend on $S_1, \ldots, S_n$, and other microenvironmental signals. With these assumptions, the corresponding term is written as:

$$R_{ik}^1 = \Omega_{ik} U_i. \tag{14}$$

- $R_{ik}^2$: Cell repair. In general, we assume that epigenetic changes, that lead to the accumulation of internal variables and to the change of cell behaviour, are reversible and that cells can repair themselves through diverse pathways. Thereupon, we include a decay term:

$$R_{ik}^2 = -\Lambda_{ik} U_i V_{ik}, \tag{15}$$

  with $\lambda_{ik}$ the decay rate.

- $R_{ik}^3$: Inheritance. Daughter cells may inherit the epigenetic changes acquired by their progenitors in a proportion $\beta_{ik} \in [0, 1]$, so that if $\beta_{ik} = 1$, daughter cells are identical to their progenitors and they have the same phenotypic state, and if $\beta_{ik} < 1$, daughter cells do not inherit all the epigenetic changes. The corresponding term is written as:

$$R_{ik}^3 = \beta_{ik} f_i U_i V_{ik}, \tag{16}$$

  where $f_i$ is the cells growth term.

- $R_{ik}^4$: Decrease due to death. When cells die, the total amount of internal variable inside a volume diminishes at rate $d_i$, so we can write:

$$R_{ik}^4 = -d_i U_i V_{ik}. \tag{17}$$

Once all the terms have been defined, we can substitute $R_{ik}$ in Eq. 13 and simplify terms, obtaining the general transport equation for internal variables $V_{ik}$:

$$\frac{\partial V_{ik}}{\partial t} = \frac{1}{U_i} \left( -\boldsymbol{q}_i \cdot \boldsymbol{\nabla} V_{ik} \right) + \Omega_{ik} - \Lambda_{ik} V_{ik} + (\beta_{ik} - 1) f_i V_{ik}, \quad i = 1, \ldots, n, k = 1, \ldots, r_i. \tag{18}$$

### B Details of parametric analyses.

#### B.1 Model parameters.

In this section we present an extended study of the parameters defining the acquisition of phenotypic changes, mediated by function $\theta$:

$$\begin{aligned} \theta &= \kappa_v(1 - \gamma F(s;k,\xi)), \\ F(s) &= \Phi(y), \\ y(s;k,\xi) &= -\frac{1}{k}\log\left(1 + \frac{s-\xi}{\xi}\right). \end{aligned}$$

First, in Figure 1 we present the effect of the different parameters on the shape of the function. Specifically, $\xi$ is a location parameter and specifies the level of oxygen below which cells undergo phenotypic changes towards a cancer stem cell phenotype. $k$ is a shape parameter defining the spread of the function and finally, $\gamma \in [1, 2]$ is a parameter setting whether cells differentiate again when the oxygen concentration is above $\xi$ , that is, if epigenetic changes are elastic/inelastic. If $\gamma = 1$, cells do not experiment any phenotypic changes when exposed to high levels of oxygen (inelastic behaviour), while if $\gamma = 2$, cells move towards differentiated phenotypes for high oxygen concentrations (elastic behaviour).

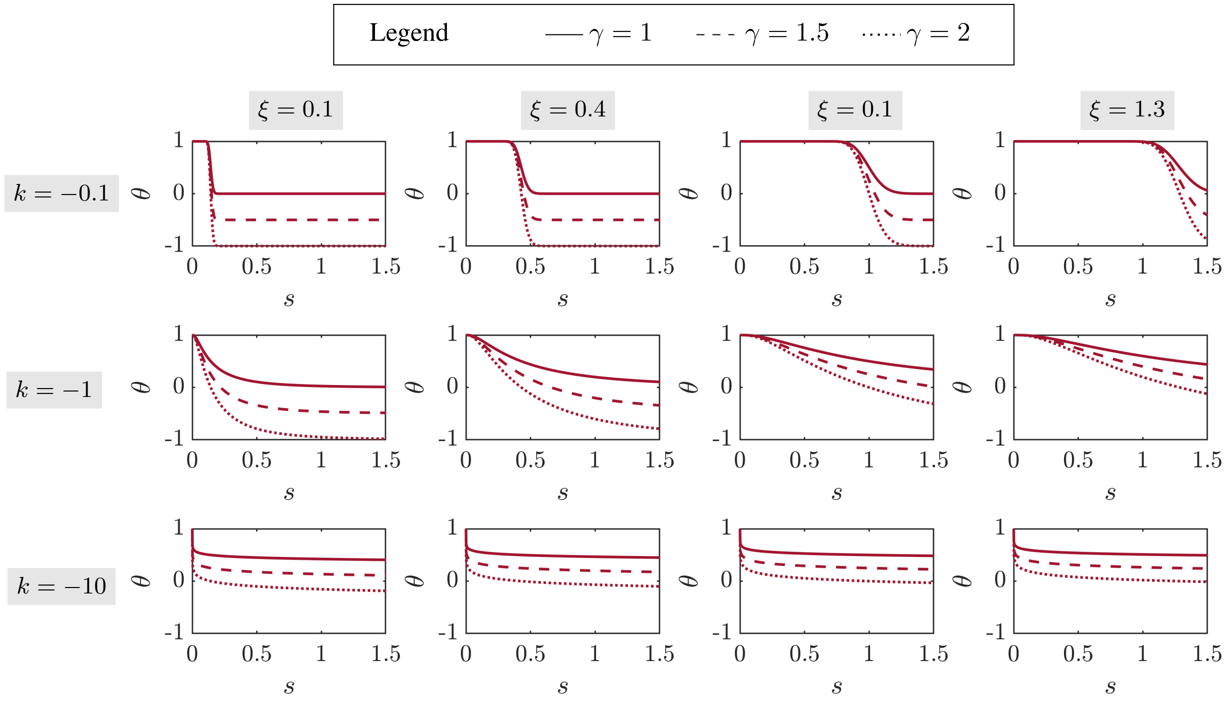


Figure 1: **Different shapes of function** $\theta$. Simulations are performed for different values of the parameters defining function $\theta$. In particular, the location parameter in function $\theta$ increases from left to right, and the shape parameter $k$ increases from bottom to top. In each subfigure, functions with different values of the repair parameter $\gamma$ are plotted with different line styles ($\gamma = 1$ with a continuous line, $\gamma = 1.5$ with a discontinuous line and $\gamma = 2$ with a dotted line).

In Figures 2, 3, 4 we show the evolution of the Tumour Burden and the Mean State for the different functions $\theta$ depicted in Figure 1 and for different initial phenotypic states.

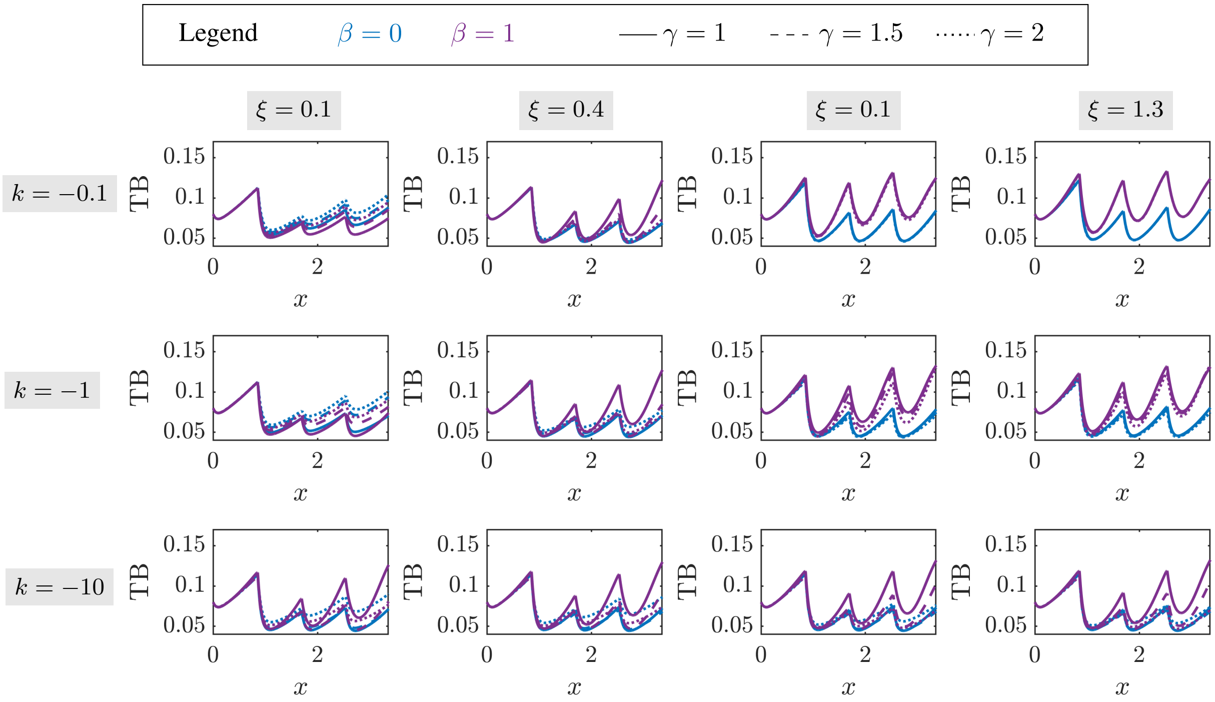


(a) Evolution of the Tumour Burden (TB).

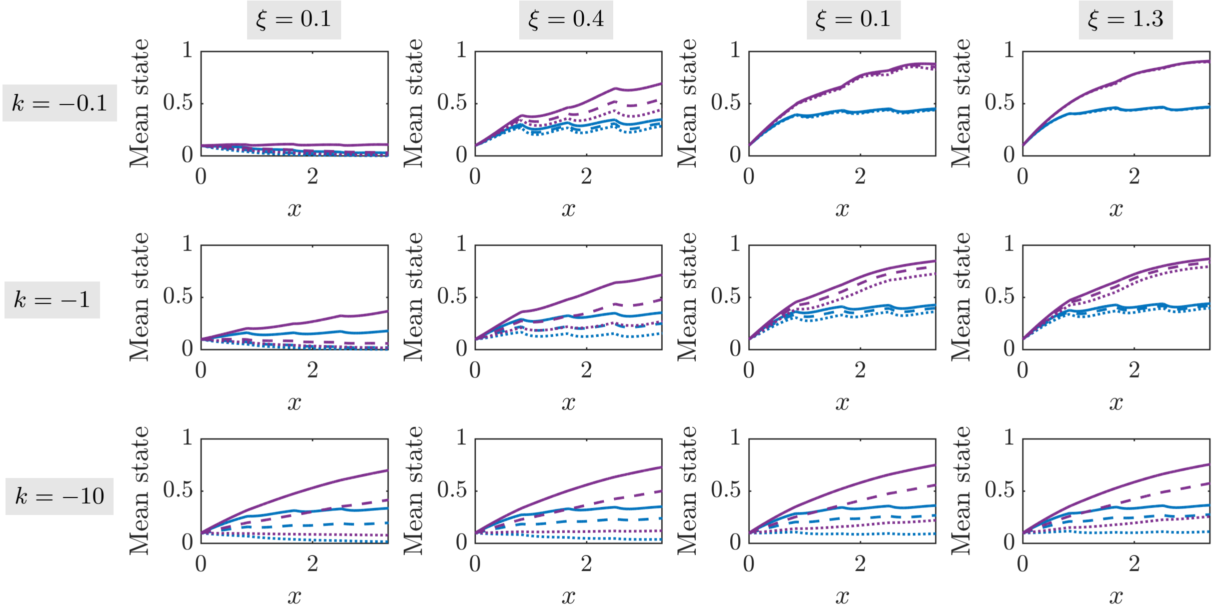


(b) Evolution of the Mean phenotypic State (MS).

Figure 2: **Evolution of the Tumour Burden (TB) and the Mean Phenotypic State for an initial phenotypic state** $v_0 = 0.1$. The grid shows simulations with different parameters related to the evolution of the phenotypic state. In particular, the location parameter in function $\omega$ increases from left to right, and the shape parameter $k$ increases from bottom to top. In each subfigure, simulations for two different $\beta$ values are plotted with different colours ($\beta = 0$ in blue and $\beta = 1$ in purple) and different values of the repair parameter $\gamma$ are plotted with different line styles ($\gamma = 1$ with a continuous line, $\gamma = 1.5$ with a discontinuous line and $\gamma = 2$ with a dotted line).

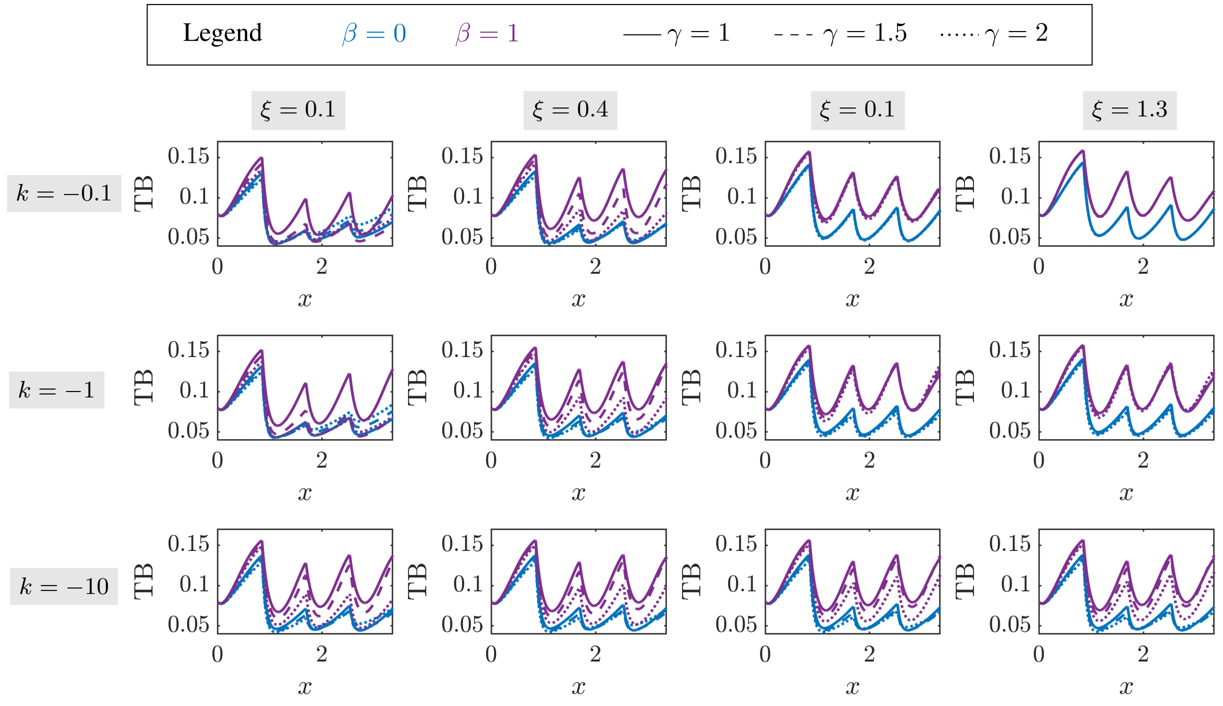


(a) Evolution of the Tumour Burden (TB).

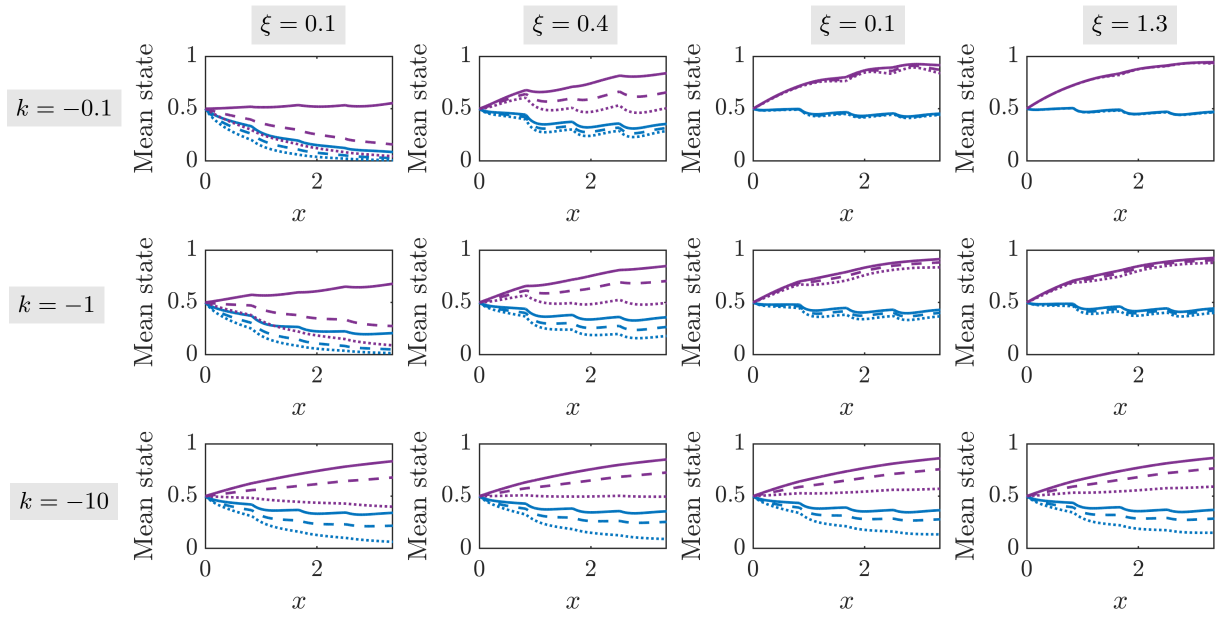


(b) Evolution of the Mean phenotypic State (MS).

Figure 3: **Evolution of the Tumour Burden (TB) and the Mean Phenotypic State for an initial phenotypic state** $v_0 = 0.5$. The grid shows simulations with different parameters related to the evolution of the phenotypic state. In particular, the location parameter in function $\omega$ increases from left to right, and the shape parameter $k$ increases from bottom to top. In each subfigure, simulations for two different $\beta$ values are plotted with different colours ($\beta = 0$ in blue and $\beta = 1$ in purple) and different values of the repair parameter $\gamma$ are plotted with different line styles ($\gamma = 1$ with a continuous line, $\gamma = 1.5$ with a discontinuous line and $\gamma = 2$ with a dotted line).

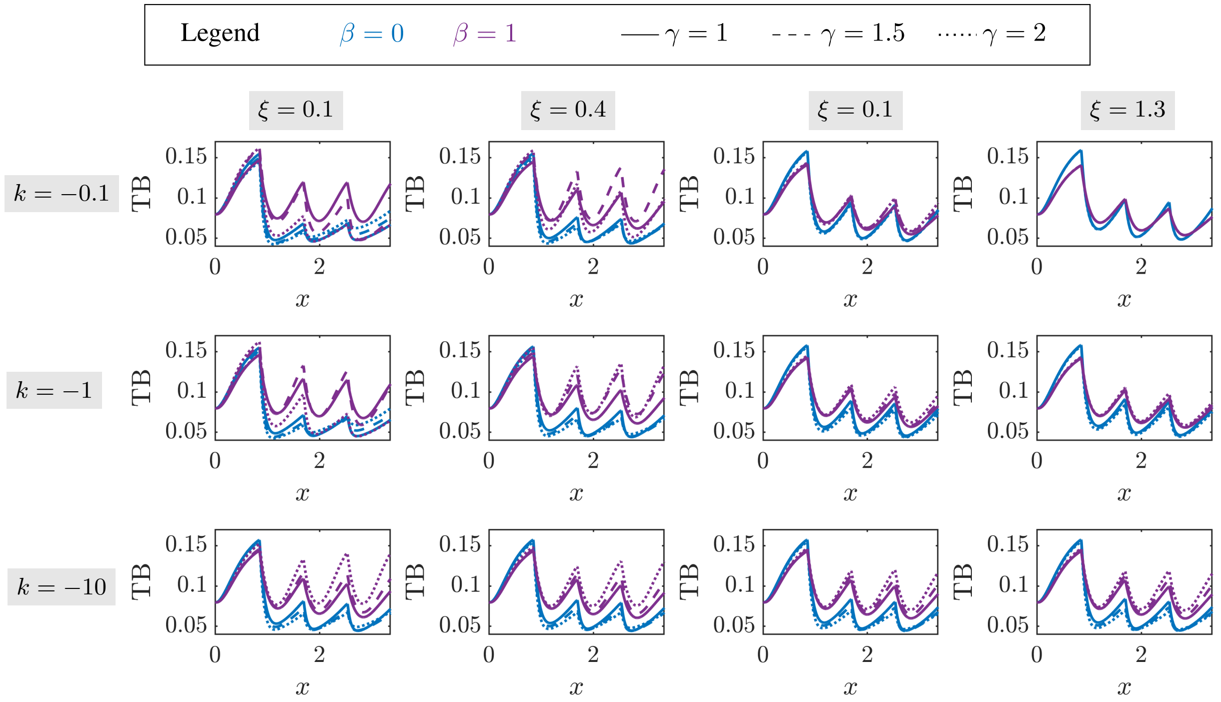


(a) Evolution of the Tumour Burden (TB).

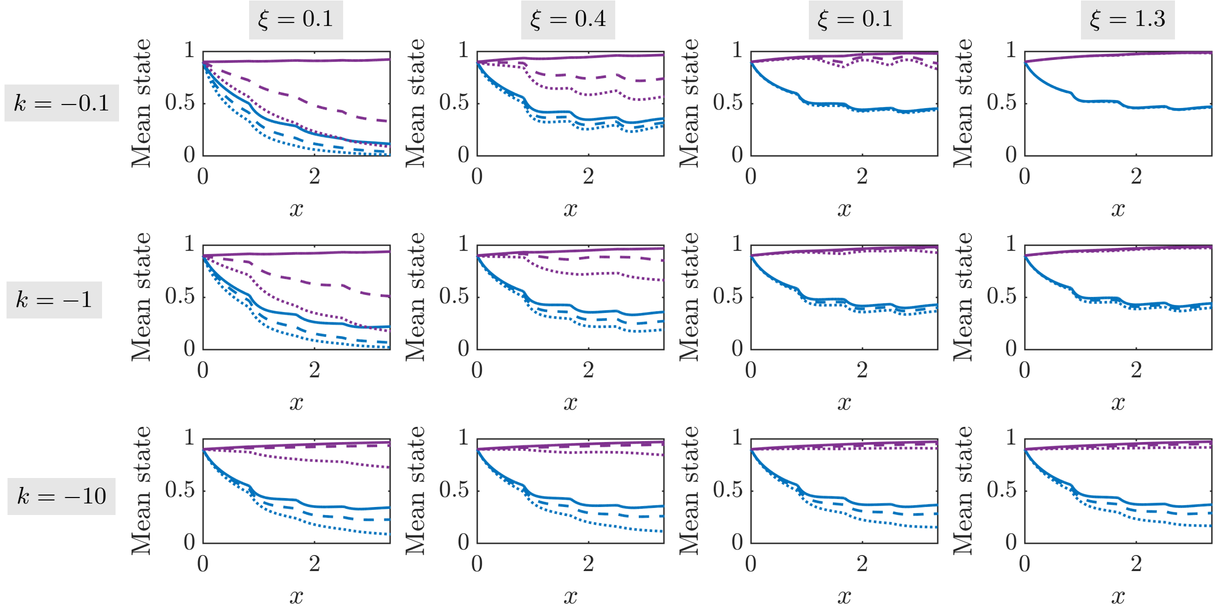


(b) Evolution of the Mean phenotypic State (MS).

Figure 4: **Evolution of the Tumour Burden (TB) and the Mean Phenotypic State for an initial phenotypic state** $v_0 = 0.9$. The grid shows simulations with different parameters related to the evolution of the phenotypic state. In particular, the location parameter in function $\omega$ increases from left to right, and the shape parameter $k$ increases from bottom to top. In each subfigure, simulations for two different $\beta$ values are plotted with different colours ($\beta = 0$ in blue and $\beta = 1$ in purple) and different values of the repair parameter $\gamma$ are plotted with different line styles ($\gamma = 1$ with a continuous line, $\gamma = 1.5$ with a discontinuous line and $\gamma = 2$ with a dotted line).

##### B.2 Environmental parameters related to the oxygenation conditions.

Here we present some illustrative results of TB evolution in different oxygenation conditions, parameterised in terms of $\mathcal{T}$, $s_{\min}$, $A$ and $\phi$. As shown by the PCCs (Figure **??**), $\mathcal{T}$ has no significant impact in the final value of the TB. Hence, we represent its variability with a band for each combination of $s_{\min}$, $A$ and $\phi$.

Table 1 presents twelve representative combinations of those three parameters, together with the observed trend in the TB (growth, regression or extinction). The temporal evolution of the TB for each of the twelve configurations mentioned in Table 1 is represented in Figure 5 with the corresponding colour (blue for TB growth, yellow for TB regression and orange for tumour extinction), together with the simulation in constant hypoxia ($s_L(t) = s_R(t) = 0.14$) and constant normoxia ($s_L(t) = s_R(t) = 1.3$).

| **Case** | $s_{\min}$ | $A$ | $\phi$ | **TB (Scenario I)** | **TB (Scenario II)** |
|---|---|---|---|---|---|
| **1** | Low (0.14) | Low (0.57) | Phase opposition ($\pi$) | Regression | Regression |
| **2** | Low (0.14) | Low (0.57) | $\pi/2$ | Extinction | Regression |
| **3** | Low (0.14) | Low (0.57) | In phase (0) | Extinction | Extinction |
| **4** | Low (0.14) | High (1.14) | Phase opposition ($\pi$) | Growth | Growth |
| **5** | Low (0.14) | High (1.14) | $\pi/2$ | Regression | Growth |
| **6** | Low (0.14) | High (1.14) | In phase (0) | Extinction | Regression |
| **7** | High (0.57) | Low (0.57) | Phase opposition ($\pi$) | Growth | Growth |
| **8** | High (0.57) | Low (0.57) | $\pi/2$ | Growth | Growth |
| **9** | High (0.57) | Low (0.57) | In phase (0) | Growth | Growth |
| **10** | High (0.57) | High (1.14) | Phase opposition ($\pi$) | Growth | Growth |
| **11** | High (0.57) | High (1.14) | $\pi/2$ | Growth | Growth |
| **12** | High (0.57) | High (1.14) | In phase (0) | Growth | Growth |

Table 1: **Summary of parameters and TB trends** Twelve representative configurations of the oxygen boundary conditions are considered, representing each possible trend.

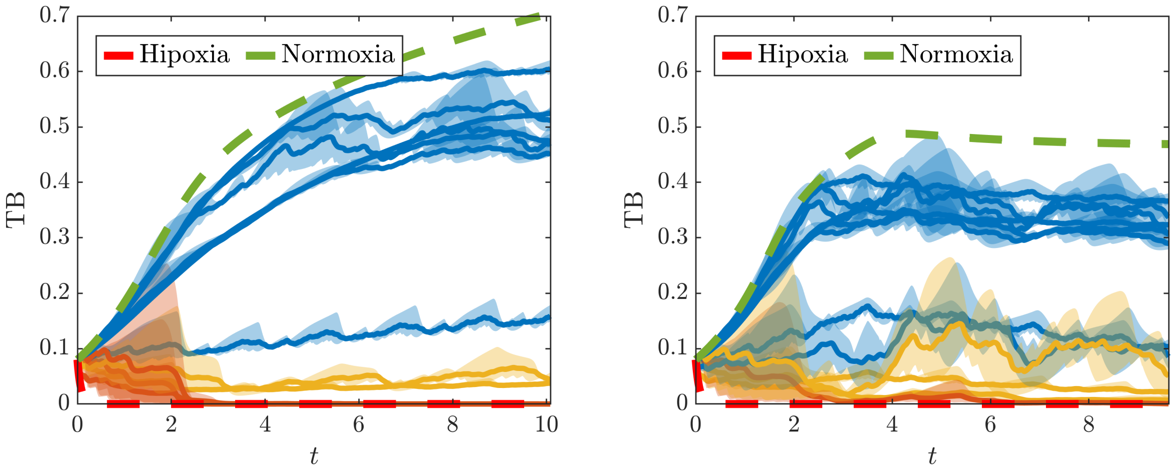


(a) Scenario I: Phenotype insensitive to oxygen variations.

(b) Scenario II: Phenotype sensitive to oxygen variations.

Figure 5: **TB evolution for the 12 cases detailed in Table 1.** A blue line indicates that the TB at the end of the simulation is bigger than at the beginning, a yellow line indicates that it is smaller, and an orange line indicates that the tumour extincts. TB evolution in both constant hypoxia ($s_L(t) = s_R(t) = 0.14$) and constant normoxia ($s_L(t) = s_R(t) = 1.3$) are depicted in red and green respectively.

Interestingly, it can be seen that, although tumours composed of cells insensitive to phenotypic variations are generally bigger at the end of the simulation, they have a higher percentage of extinction. This suggests that phenotypic plasticity endows the tumours with a kind of resilience, even if that means being smaller in general, at least in what regards the number of alive cells.
